# Amino acid substitutomics: profiling amino acid substitutions at proteomic scale unveils biological implication and escape mechanism in cancer

**DOI:** 10.1101/2024.11.18.624056

**Authors:** Peize Zhao, Shengzhi Lai, Shuaijian Dai, Chen Zhou, Ning Li, Weichuan Yu

**Affiliations:** Individual Interdisciplinary Program, The Hong Kong University of Science and Technology, Hong Kong, China; Department of Electronic and Computer Engineering, The Hong Kong University of Science and Technology, Hong Kong, China; Division of Life Science, The Hong Kong University of Science and Technology, Hong Kong, China; Department of Biomedical Engineering, School of Basic Medical Sciences, Central South University, Hunan Province, China

**Keywords:** Cancer proteome, Biomarker, Cancer escape, Amino acid substitutomics, Amino acid substitutome

## Abstract

Amino acid (AA) substitutions play a critical role in regulating cellular activities, including complex signaling and cell cycle processes. Recent research on AA substitutions has primarily relied on genomic and transcriptomic data. The identification at the proteomic scale remains underexplored, despite evidence suggesting that DNA and RNA biosynthesis are not the sole sources of these substitutions. This gap persists due to challenges in analyzing large-scale proteomic data. In this study, we address this limitation by analyzing multiple independent datasets across five cancer types using PIPI-C, a novel mass spectrometry data analysis tool. And we propose AA substitutomics, a pipeline for characterizing AA substitutions arising after protein translation and dissecting the regulatory functions of key proteins with AA substitutions. Among our identified AA substitutions, 87% are novel findings and not recorded in genomic/transcriptomic databases, which indicates that the post-translational AA substitutions are prevalent. Our findings reveal biologically significant AA substitutions linked to cancer, such as F43S and E91D in hemoglobin subunit beta, P584T in filamin A, and A175N in fructose-bisphosphate aldolase B. Furthermore, our pipeline enables direct investigation of drug resistance and immune escape. By capturing functional protein-level alterations beyond genomic and transcriptomic profiling, it establishes a robust framework to advance cancer research.

## 1 INTRODUCTION

Cancer is a heterogeneous group of malignancies defined by uncontrolled cellular proliferation, invasion, and metastasis, driven by the breakdown of intrinsic cellular regulatory networks that govern normal growth and homeostasis [1]. As the primary functional executors of nearly all cellular processes, proteins play an indispensable role in tumorigenesis: aberrant protein expression, mutational damage, or functional dysregulation directly disrupts core cellular behaviors including division, survival, and apoptosis, forming the molecular basis of malignant transformation [2, 3]. While dysregulated oncogenic and tumor-suppressive signaling pathways are well-established drivers of cancer, the role of proteome-level alterations—particularly amino acid (AA) substitutions—remains incompletely defined, despite their direct impact on protein structure and function.

A critical yet incompletely characterized driver of cancer progression is the accumulation of AA substitutions within cellular proteomes. These alterations disrupt native protein structure and function, with a subset conferring selective growth advantages to tumor cells and reinforcing core malignant hallmarks [4, 5, 6]. AA substitutions arise from three biologically distinct, non-overlapping sources: DNA mutations, mRNA mutations, and errors during protein translation [7]. Notably, the relative contribution of each source to the total pool of cancer-associated AA substitutions remains poorly defined, representing a critical gap in current understanding of tumor proteomic dysregulation (Figure 1A). Research into cancer-associated AA substitutions has been revolutionized by multi-omics integration, with each omics layer contributing unique insights into tumor pathogenesis and progression. Genomics forms the foundational framework, enabling identification of driver DNA mutations, chromosomal aberrations, and copy number variations that initiate oncogenic transformation [8, 9]; prior genomic analyses indicate a substantial proportion of AA substitutions are deleterious, with proteins carrying these harmful variants frequently localized to pivotal nodes in protein-protein interaction networks, though precise quantification of this detrimental subset requires cross-validation with modern datasets [9]. Building on this genomic foundation, transcriptomics profiling delineates global gene expression patterns, yielding actionable prognostic and predictive biomarkers, defining tumor heterogeneity, and uncovering molecular drivers of oncogenic progression [10]. Key transcriptomic advances include the identification of cancer-specific gene signatures and epitranscriptomic alterations; for instance, RNA editing of podocalyxin-like in the gastric cancer cell line reduces cellular proliferation and invasiveness compared to the unedited transcript variant [11]. While genomic and transcriptomic approaches have dominated cancer substitution research, emerging proteomic studies bridge the critical gap between genotype and functional phenotype. For example, a large-scale analysis of 1,823 tumor samples across 11 cancer types detected significant enrichment of tryptophan-to-phenylalanine (W>F) substitutions, with liver tumors showing a four-fold higher prevalence of this alteration relative to the pan-cancer average [12]. These W>F substitutions, linked to non-canonical translational decoding mechanisms, disrupt target gene function and modulate tumor immune surveillance. Additionally, arginine-to-cysteine (R>C) substitutions are highly enriched in Kelch-like ECH-associated protein 1-mutant lung cancers, with 33.5% of R>C-containing peptides shared between two distinct lung cancer subtypes, indicating conserved substitution patterns across histologically related malignancies [13].

**Figure 1:**
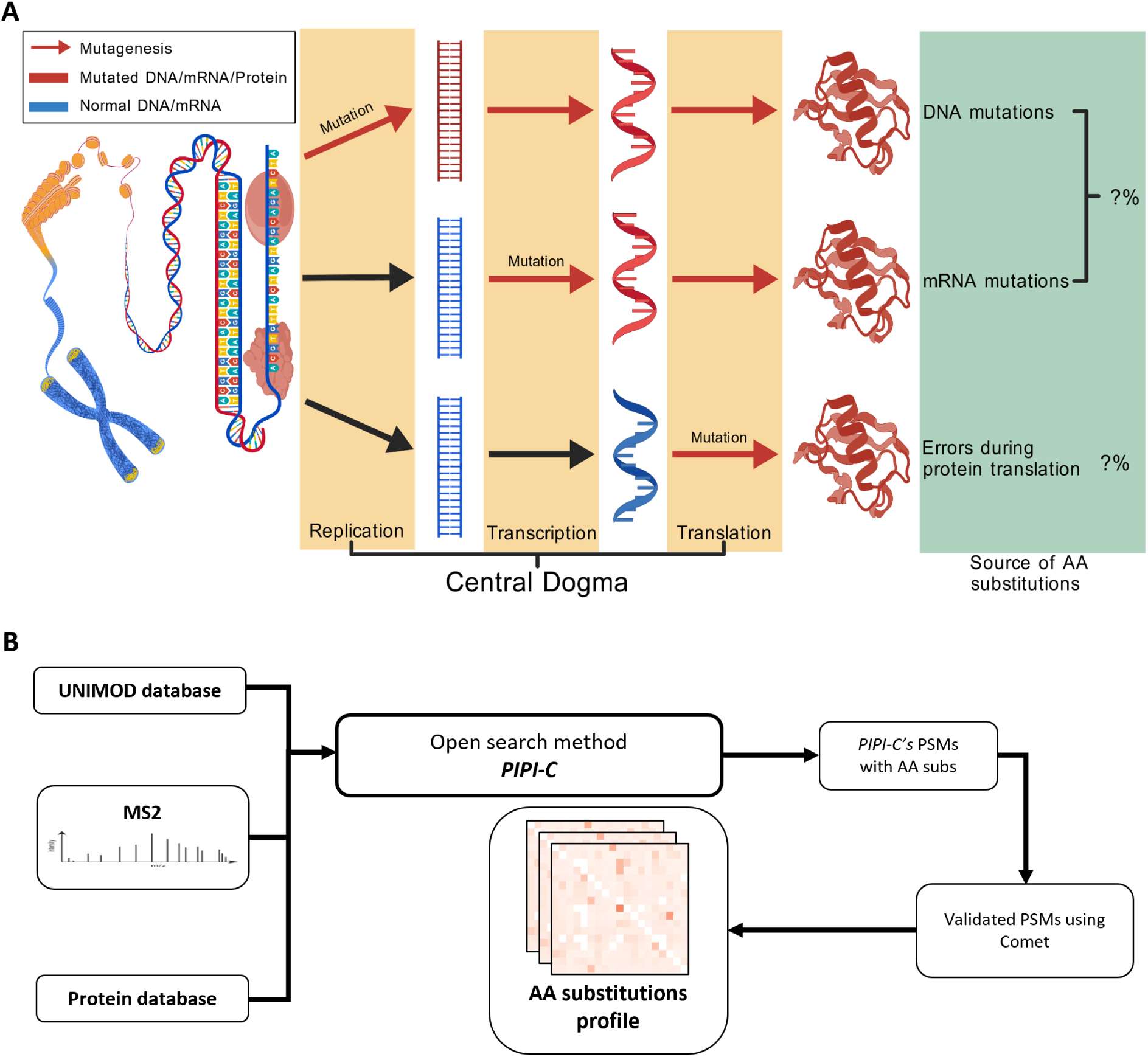
Demonstration of AA substitutomics. (A) The diagram illustrates three distinct sources of AA substitutions. AA substitutions can arise independently from misincorporation events occurring during DNA replication, mRNA synthesis, and protein translation, respectively. In principle, AA substitutomics refers to the global profiling of all AA substitutions present within a cell. In this study, we specifically focus on AA substitutions originating from translational errors and intracellular post-translational biochemical modifications of proteins. (B) The workflow of AA substitutomic analysis.

Despite the established biological relevance of AA substitutions in cancer proteomics, comprehensive and unbiased identification of these alterations at the global proteomic scale remains a substantial technical challenge. Existing studies have been limited to targeted analyses of a narrow range of AA substitutions, leaving the complex interplay between multi-site substitutions, their regulatory networks, and cancer-type specificity poorly resolved across diverse malignancies. Traditional proteomic database search strategies rely on closed-search workflows that only detect predefined post-translational modifications (PTMs) with fixed modification parameters, restricting discovery of unannotated or novel substitutions [14, 15]. While these closed-search approaches deliver high accuracy for targeted modifications, they suffer from exponential database expansion and reduced sensitivity when assessing multiple concurrent modifications, limiting their utility for global substitution profiling. This limitation has driven the development of open-search methodologies, which enable unbiased interrogation of large modification databases such as UNIMOD to identify uncharacterized PTMs and AA substitutions [16]; historically, however, open-search strategies have traded search breadth for identification precision, hindering their broad clinical and research application. Recent technical advances in open-search algorithms have refined this balance, optimizing both search space coverage and identification accuracy to enable systematic, global profiling of all AA substitution types from complex proteomic datasets [17, 18, 19]. Among these advanced tools, PIPI-C stands out for its combinatorial optimization framework for peptide identification and PTM characterization, leveraging fuzzy bidirectional matching to map short peptide sequences within protein structures and solving a mixed integer linear programming model to accurately determine peptide sequences and their corresponding PTM patterns [17].

In this study, we implement a dual-search strategy that integrates the broad discovery capacity of open-search with the high precision of closed-search, using PIPI-C as the primary open-search engine and the widely validated Comet algorithm as the closed-search validation tool (Figure 1B). To ensure findings are robust and generalizable across diverse tumor types, we apply this optimized analytical pipeline to multiple proteomic datasets covering five distinct cancer types, minimizing tumor-specific biases and technical variability inherent in single-cohort analyses. Furthermore, we formally define and establish the novel concept of AA substitutomics, a specialized analytical framework focused on characterizing AA substitutions derived from translational errors, as well as dissecting the regulatory functions of key proteins carrying multiple high-significance AA substitutions. This AA substitutomics pipeline uncovers cancer-specific substitution signatures that correlate with distinct protein functional alterations, particularly within core oncogenic signaling pathways and protein interaction networks. To validate the functional relevance of key identified substitutions, we integrate a comprehensive review of proteomic and protein structural literature to enhance result reliability. Beyond mechanistic characterization, this novel pipeline offers a unique approach to exploring the molecular mechanisms underlying cancer drug resistance and immune evasion driven by proteomic substitutions. Ultimately, the AA substitutomics framework provides critical insights into the fundamental molecular basis of tumor biology, clarifying how specific AA modifications shape cancer phenotypes by altering protein stability, intermolecular binding affinity, and downstream functional activity.

## 2 RESULTS

### 2.1 AA substitutions landscape across five cancer cohorts

PIPI-C is an open-search peptide identification platform built on a mixed integer linear programming (MILP) framework, enabling efficient simultaneous characterization of diverse PTMs and AA substitutions [17]. Detailed information regarding the algorithmic design and practical implementation of PIPI-C is provided in Supplementary Note 1. To establish a robust analytical pipeline for global AA substitution profiling, we systematically compared and validated the identification performance of PIPI-C against two cutting-edge open-search tools, Open-pFind [18] and MODplus [19], using a multi-layered validation framework. Our rigorous validation workflow comprised three core modules: (1) assessment of false positive rates using simulated and synthetic peptide datasets devoid of AA substitutions (Supplementary Note 2); (2) evaluation of detection sensitivity using simulated datasets containing 1–3 predefined AA substitutions per peptide (Supplementary Note 3); and (3) verification of multi-modification identification capacity using experimental proteomic data with serially enriched phosphorylation, ubiquitination, and acetylation modifications [20] (Supplementary Note 4).

In the false positive validation assay, PIPI-C exhibited a low false identification rate of up to 0.3% in substitution-free simulated datasets (Supplementary Figure S2) and approximately 1% in substitution-free synthetic peptide datasets (Supplementary Figure S3), confirming its high specificity. For sensitivity testing, we employed three distinct simulated datasets, each containing 5% of spectra harboring one, two, or three AA substitutions per peptide, respectively. In the single AA substitution scenario, Open-pFind achieved a detection ratio closer to the theoretical 5% threshold relative to the other two tools. However, under conditions of multiple concurrent AA substitutions per peptide, PIPI-C outperformed both Open-pFind and MODplus, with a detection ratio consistently approaching the 5% baseline. In multi-modification identification assays, PIPI-C also demonstrated superior performance, particularly for peptides carrying multiple distinct PTMs (Supplementary Figure S4). As a representative example, PIPI-C identified 2,604 peptide spectrum matches (PSMs) containing two or more acetylation sites per peptide, markedly outperforming Open-pFind (609 PSMs) and MODplus (1,256 PSMs). Collectively, these comparative analyses confirm the superior performance of PIPI-C in detecting peptides with multiple co-occurring modifications, a critical technical requirement for unbiased AA substitutomics analysis. We further validated the capacity of PIPI-C to distinguish biologically relevant, enriched AA substitutions from non-specific, non-enriched substitutions (Supplementary Note 5), further verifying its robustness and reliability for the analytical pipeline employed in this study. These rigorous validations collectively establish PIPI-C as the optimal tool for subsequent AA substitutomics profiling.

Previous studies have characterized cancer-associated protein molecular dysregulation and tumorspecific AA substitution signatures linked to oncogenic pathways [12, 13], yet these investigations were restricted to targeted analysis of a limited subset of AA substitutions, lacking pan-cancer, unbiased profiling of all substitution types. To address this gap, the present study performed a comprehensive, unbiased analysis of global AA substitution profiles across five distinct cancer types, leveraging high-quality proteomic data from the Clinical Proteomic Tumor Analysis Consortium (CPTAC) cohorts [21]. The integrated dataset encompasses a total of 537 patients across five cancer types, including 99 cases of glioblastoma multiforme (GBM) [22], 110 cases of head and neck squamous cell carcinoma (HNSCC) [23], 110 cases of lung adenocarcinoma (LUAD) [24], 108 cases of lung squamous cell carcinoma (LSCC) [25], and 110 cases of clear cell renal cell carcinoma (ccRCC) [26]. These five cancer types were selected based on three stringent criteria to ensure data reliability and result generalizability: first, comparable cohort sizes (100–110 patients per cancer type) to minimize sample size-related bias; second, uniform proteomic acquisition protocols utilizing isobaric TMT-10/11 labeling, with consistent reporter ion mass shifts (229.163 Da) to guarantee quantitative accuracy and comparability; third, histopathological and anatomical diversity, covering neuroectodermal tumors, adenocarcinomas, and squamous cell carcinomas derived from distinct organ systems. This study design balances technical uniformity and histopathological diversity, enhancing the representativeness and broad applicability of the resulting findings.

The overall proteomic data analysis workflow is outlined in Figure 1B. Briefly, all mass spectrometry spectra from the five CPTAC cancer datasets [21] and all modification entries in the UNIMOD database [16] were used as input for PIPI-C-based open-search analysis. AA substitution identified by PIPI-C were subsequently validated using the Comet closed-search algorithm, with only crossvalidated results retained for downstream profiling and analysis. Both PIPI-C and Comet were run with a PSM-level false discovery rate (FDR) cutoff of 1% to ensure result quality. We also identified potential identification ambiguities stemming from identical mass shifts between specific AA substitutions and other PTMs (Supplementary Table S4), and all such ambiguous substitutions were excluded from subsequent analyses to eliminate false positive calls. Detailed search results of the AA substitutomics pipeline and cross-cohort generalization test data are presented in Supplementary Figure S7.

Normalized ratios of individual AA substitution types across the five cancer cohorts are visualized as heatmaps in Figure 2A, revealing several conserved pan-cancer substitution patterns. Notably, among all glutamic acid derived substitutions (E>X), the E>M substitution was the most abundant, accounting for 23–28% of all E-related substitutions across cohorts. Similarly, among proline derived substitutions (P>X), the P>V substitution was the most prevalent, representing 25–38% of total P-related substitutions (Supplementary File 1). Both raw PIPI-C identification results and Comet-validated AA substitution datasets are publicly available on Zenodo [27] under record number 13383150, ensuring full data reproducibility.

**Figure 2:**
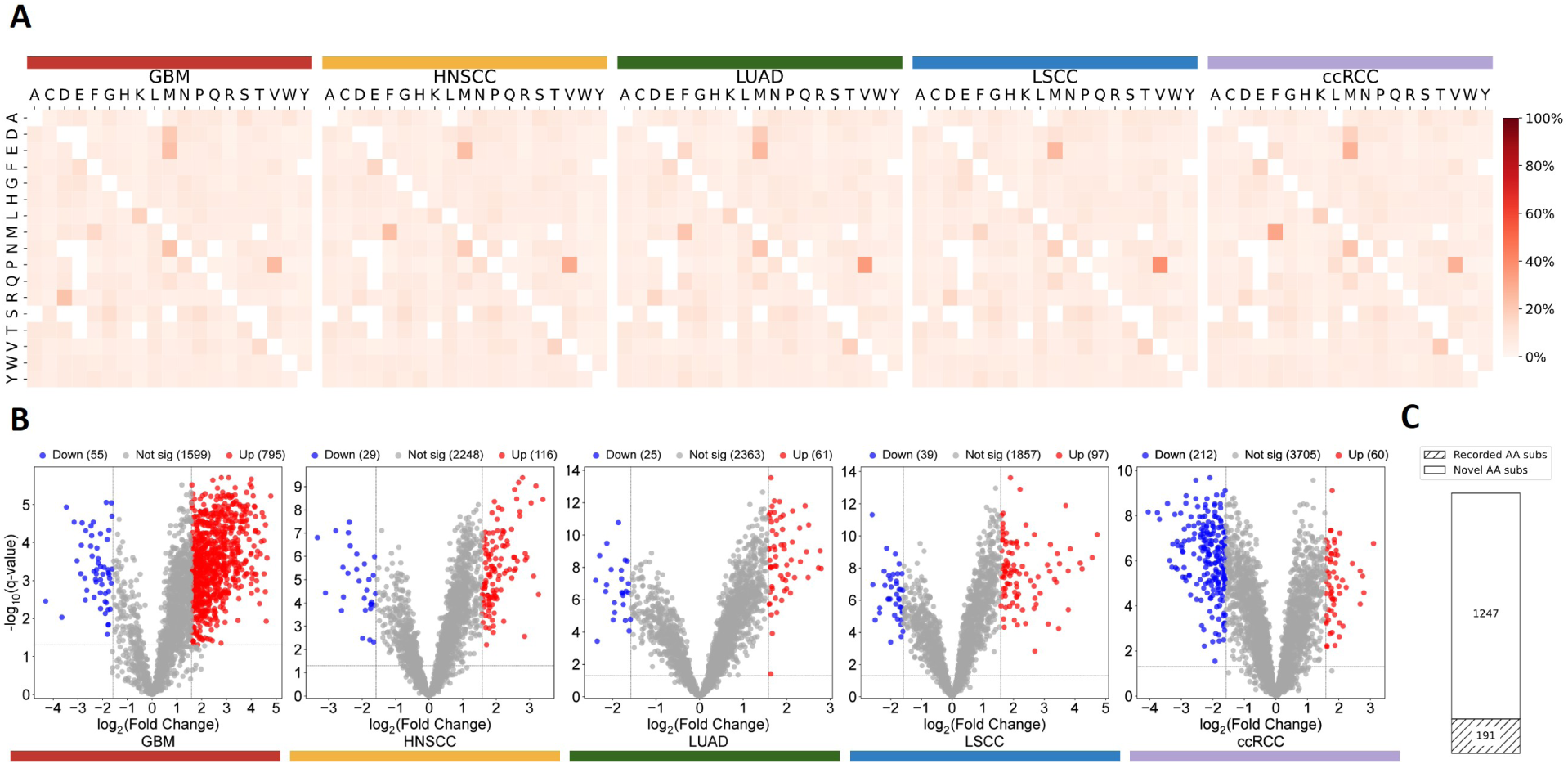
AA substitutions landscape across five cancer cohorts. (A) Heatmaps show the normalized ratios of all types of AA subs in validated PSMs among five cancer cohorts. The horizontal AAs are the initial AAs, and the vertical AAs are the AAs after substitution. The shade of red represents the normalized ratio of AA substitutions. The color bars represent five cancer cohorts: GBM (dark red), HNSCC (yellow), LUAD (green), LSCC (blue), and ccRCC (purple) (Supplementary File 1). For the search results of the AA substitutomic pipeline, please refer to Supplementary Figure S7. (B) Volcano plots of quantification results based on the validated results. Down-regulated UPSPs are denoted by blue. Up-regulated UPSPs are denoted by red. Not significantly regulated UPSPs are denoted by gray (Supplementary File 2). The color bars represent five cancer cohorts: GBM (dark red), HNSCC (yellow), LUAD (green), LSCC (blue), and ccRCC (purple). (C) Bar plot showing 191 out of 1438 identified AA substitutions are recorded in the protein variant database (Supplementary File 3). Clean section: Novel AA substitutions. Slashed section: Recorded AA substitutions.

We next performed quantitative profiling to assess differential regulation of AA substitutions containing unique PTM site patterns (UPSPs) between tumor and matched normal tissue samples (Figure 2B, Supplementary File 2). For comprehensive reference, differential regulation profiles of UPSPs carrying all modification types are shown in Supplementary Figure S8. A strict foldchange threshold was applied for significance calling: UPSPs with a fold change > 3 were defined as up-regulated, while those with a fold change *<* 1/3 were categorized as down-regulated; detailed quantitative methodologies and threshold justification are described in the Methods section. Quantitative analysis revealed that the majority of AA substitution-modified UPSPs were not significantly differentially regulated between tumor and normal samples. Among significantly regulated UPSPs, only 13% (191 out of 1438) of the identified AA substitutions were annotated in the UniProt protein variant database [28], while the remaining 1247 AA substitutions represent novel, uncharacterized variants with potential biological relevance in cancer biology (Figure 2C, Supplementary File 3).

Notably, AA substitutions were unevenly distributed across the proteome in all cancer cohorts. Hemoglobin subunit beta (HBB) was identified as one of the proteins with the highest number of AA substitutions, with a total of 104 distinct substitutions detected across the five cohorts, despite its short amino acid length of only 147 residues (Supplementary Figure S9); consistent with pan-cancer patterns, the E>M substitution remained the most abundant substitution in HBB. Additional proteins harboring multiple significantly regulated AA substitutions are listed in Supplementary Figure S10. Differential regulation patterns varied markedly across cancer types: in four cohorts (GBM, HNSCC, LUAD, and LSCC), the number of up-regulated UPSPs exceeded that of downregulated UPSPs, with GBM exhibiting the highest number of up-regulated UPSPs (795). In stark contrast, ccRCC displayed an inverse regulatory pattern, with significantly more down-regulated UPSPs (212) than up-regulated UPSPs (60). These distinct cohort-specific quantitative profiles highlight pronounced cancer-type-specific AA substitution regulation, which we further interrogated in subsequent focused analyses of significantly regulated UPSPs.

### 2.2 Multiple significantly regulated AA substitutions as cancer biomarkers

Based on the findings of our AA substitutomic analysis pipeline, we identified eight cancer biomarkers linked to multiple significantly regulated AA substitutions. To elucidate the biological implications of these crucial proteins and the associated AA substitutions, we performed GO and KEGG pathway enrichment analyses [29, 30]. Comprehensive descriptions of the AA substitutions in proteins related to extracellular exosomes, actin filament binding, and the glycolysis/gluconeogenesis pathways are provided in the subsequent sections.

#### 2.2.1 AA substitutomes in extracellular exosome proteins across cancer cohorts

Extracellular exosomes, also known as microvesicles, are a type of cell-excreted extracellular vesicle that is normally less than 100 nanometers in diameter. As the smallest extracellular vesicle, exosomes are crucial in the regulation of physiological processes including immune response, neuronal communication, and cancer development. Because of the endogenous nature of exosomes, they exhibit enhanced biocompatibility and stability when compared to synthetic carriers, making them valuable as delivery vehicles between cells [31]. Some researchers have articulated the relationship between microvesicles and cancers. In the cancer microenvironment, exosomes can take the role of extracellular messengers. For example, exosomes derived from GBM cells have been documented to trigger glioma cell proliferation and enhance cancer growth in a self-sustaining fashion [32]. Elevated exosome release stands out as a critical phenotype of malignant tumors when regulated by tumor extracellular acidity [33].

In this study, GO enrichment analysis revealed that the term extracellular exosome was significantly enriched across all five cohorts (Supplementary Figure S11). Among cellular component terms showing prominent enrichment in at least four cohorts, four of the top eight were associated with the extracellular compartment (Figure 3A), suggesting a potential association between AA substitutions and the tumor extracellular microenvironment. The counts of exosome-related genes per cancer type are displayed in Figure 3B. GBM exhibits the highest number of upregulated AA substitutions in exosome-related proteins (86), exceeding the other four cancer types. The number of exclusively upregulated proteins is 86, 32, 36, 43, and 11 in GBM, HNSCC, LUAD, LSCC, and ccRCC, respectively.

**Figure 3:**
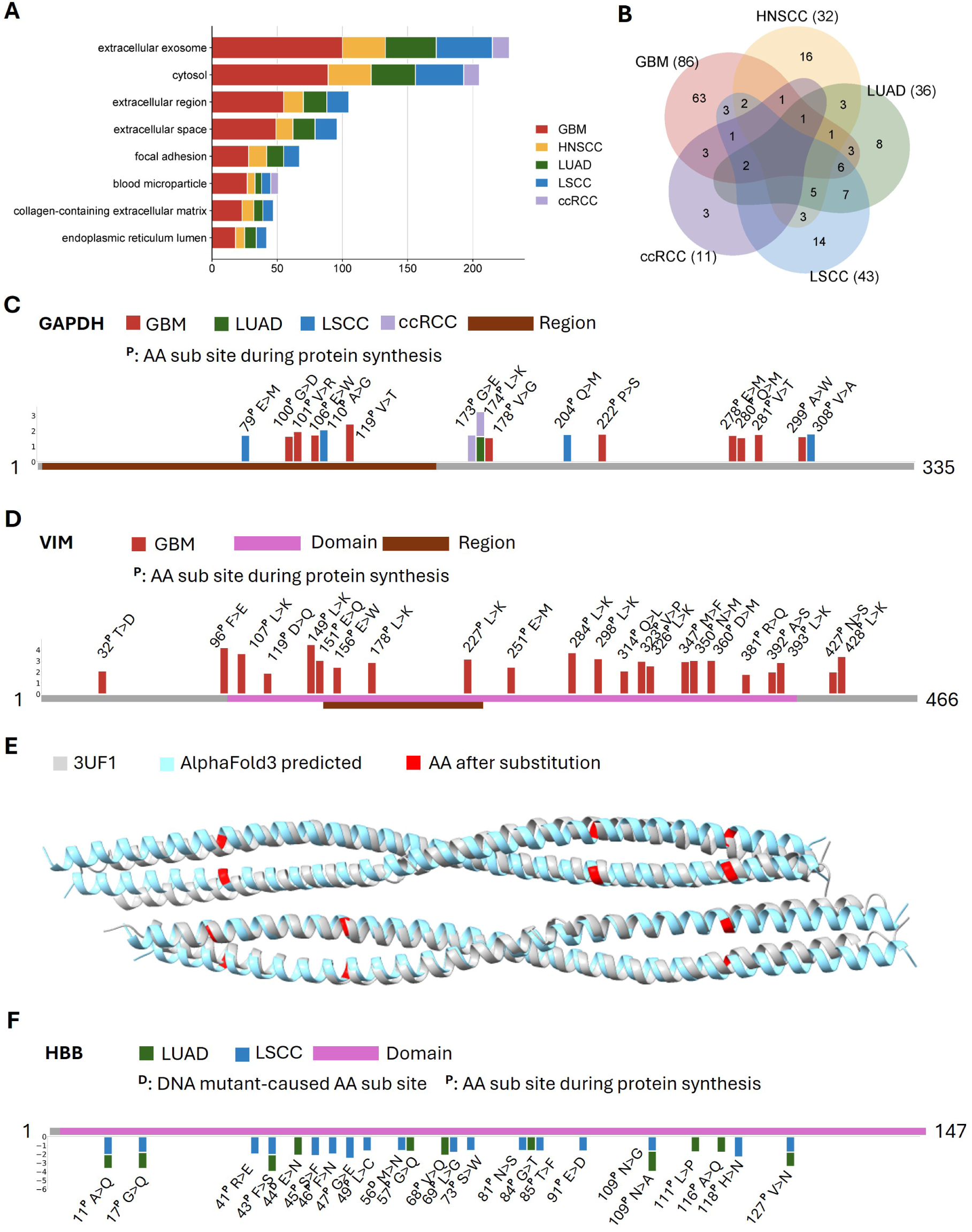
AA substitutomes in extracellular exosome proteins across cancer cohorts. (A) Bar plot showing the cellular component terms of GO enrichment that are expressed in at least four cancer cohorts. The cohorts are GBM (dark red), HNSCC (yellow), LUAD (green), LSCC (blue), and ccRCC (purple). (B) Protein overlap of only up-regulated genes. The numbers of only up-regulated genes in each cohort are noted. The cohorts are GBM (dark red), HNSCC (yellow), LUAD (green), LSCC (blue), and ccRCC (purple). (C) Cumulative AA substitutions map of GAPDH across different cancer cohorts. The AA substitutions above the protein sequence are up-regulated. Barplot showing the base 2 logarithm ratio of cancer regulation (Supplementary File 4). Brown segment: binding and interaction region. ^P^: AA substitution site during protein synthesis. The cohorts are GBM (dark red), LUAD (green), LSCC (blue), and ccRCC (purple). (D) Cumulative AA substitutions map of VIM in GBM. The AA substitutions above the protein sequence are up-regulated. Barplot shows the base 2 logarithm ratio of cancer regulation (Supplementary File 4). Brown segment: binding and interaction region. Pink segment: structural domain. ^P^: AA substitution site during protein synthesis. The cohort is GBM (dark red). (E) Structure comparison between the X-ray diffraction structure of human vimentin (fragment 144–251) (PDB: 3UF1) and structure prediction with cumulative AA substitutions. Gray: the structure of 3UF1. Light blue: AlphaFold3-predicted structure with cumulative AA substitutions in fragments 144–251. Red: The AAs after substitution. (F) Cumulative AA substitutions map of HBB in two types of lung cancers. The AA substitutions below the protein sequence are down-regulated. Barplot showing the base 2 logarithm ratio of cancer regulation (Supplementary File 4). Pink segment: structural domain. ^D^: DNA mutant-caused AA substitution site. ^P^: AA substitution site during protein synthesis. The cohorts are LUAD (green) and LSCC (blue).

We characterized the detailed AA substitution profiles of three key exosome-associated proteins: glyceraldehyde 3-phosphate dehydrogenase (GAPDH), vimentin (VIM), and HBB (Figure 3C, D and F). For GAPDH, we mapped ten AA substitutions in GBM, one in LUAD, four in LSCC, and two in ccRCC (Figure 3C, Supplementary File 2). A functional interaction region spanning residues 2 to 148 in GAPDH mediates binding with tryptophanyl-tRNA synthetase 1 (WARS1), a critical enzyme regulating aminoacylation during protein translation [34]. Within this region, we identified four upregulated AA substitutions in GBM (sites 100, 101, 106, 119) and two upregulated substitutions in LSCC (sites 70, 110). These substitutions are predicted to disrupt the GAPDH-WARS1 interaction and impair downstream aminoacylation activity, altering global protein synthesis in tumor cells.

For VIM, a total of 23 substitution sites were identified as downregulated (Figure 3D), while two distinct upregulated AA substitutions were detected at site 251: E251M (16-fold change) and E251F (5.54-fold change) (Supplementary File 2). The intermediate filament (IF) rod domain, spanning residues 103 to 411, constitutes 66% of the full VIM sequence and harbors 22 of the identified substitution sites; this domain forms the core structural scaffold of VIM and is essential for cytoskeletal microtubule assembly and cellular structural integrity [35]. Within this rod domain, a coiled-coil region spans residues 154 to 245. To evaluate the structural impact of AA substitutions on this region, we utilized AlphaFold3 to predict the structure of the modified sequence spanning residues 144 to 251 (carrying the E251M substitution, given its higher fold change) and compared it to the corresponding wild-type X-ray diffraction structure [36] (Figure 3E). Six upregulated AA substitutions fall within this sequence range, and cumulative structural alterations are predicted to destabilize microtubule-associated exosomes, potentially promoting GBM tumor cell invasion and metastasis.

Notably, the two lung cancer cohorts (LUAD and LSCC) exhibit significant downregulation of AA substitutions specifically in HBB. In LUAD, 19 of 23 significantly downregulated UPSPs map to HBB, while in LSCC, 14 of 38 downregulated UPSPs are derived from HBB. Additionally, seven UPSPs from hemoglobin subunit alpha 1 (HBA1) are significantly downregulated in LSCC; neither HBB nor HBA1 show similar downregulation in the other three cancer types. These two proteins are the primary subunits of hemoglobin, responsible for pulmonary oxygen transport to peripheral tissues, and their coordinated downregulation of AA substitutions in lung cancers suggests a disease-specific functional signature. This finding may provide novel proteomic-based strategies for lung cancer diagnosis and targeted therapy. HBB carries one of the highest densities of identified AA substitutions, all of which are downregulated and exclusive to the two lung cancer cohorts. All detected HBB substitutions localize to its globin domain (residues 3 to 147) (Figure 3F), with 17 substitution sites identified in LSCC and 11 in LUAD. Among these, sites 44 and 56 correspond to known DNA mutation-driven AA substitutions (E44K and M56I, respectively) documented in previous studies [37] (Supplementary File 2). Two substitution sites in LSCC carry two distinct variants: site 69 harbors L69G (0.28-fold) and L69Q (0.30-fold), while site 118 carries H118N (0.20fold) and H118E (0.25-fold). Several downregulated substitutions are shared between LUAD and LSCC, including A11Q, G17Q, and F43S, which may represent conserved molecular mechanisms underlying lung cancer pathogenesis.

We identified two well-validated AA substitution sites in HBB through comprehensive literature review. The F43S substitution has been shown via peptide fingerprint analysis to alter the heme pocket configuration, leading to severe hemolytic anemia [38, 39]. The F43 residue in HBB is evolutionarily conserved and plays a crucial role in maintaining hemoglobin structural integrity via its hydrophobic properties. Substitution with a polar serine residue induces significant conformational changes, destabilizing the hemoglobin quaternary structure. This phenylalanine residue is strategically positioned adjacent to the heme group’s porphyrin ring, facilitating optimal planar orientation and strong van der Waals interactions critical for molecular stability [39]. The introduction of a polar serine residue disrupts these favorable van der Waals interactions and weakens two key intra-chain hydrogen bonds (F43–G47 and F43–T39), further destabilizing the protein architecture. Given the porphyrin ring’s essential role in oxygen binding [40], this substitution may profoundly impair oxygen transport efficiency, with critical implications for the hypoxic tumor microenvironment in lung cancer patients.

Another well-characterized AA substitution in HBB is E91D, which has been shown to enhance oxygen-carrying efficiency via liquid chromatography (LC) profiling [41]. Although residue 91 is surface-exposed and does not directly interact with the heme pocket, it is located proximal to the heme-bound H93 and adjacent to two hydrophobic residues (L89, L92) within the heme-binding pocket. Unlike the reduced oxygen affinity associated with E91L and E91G substitutions, E91D confers increased oxygen affinity. Previous studies suggest that E91 elevates the pKa value of H147 in the deoxygenated state, stabilizing the protein’s quaternary structure and enhancing oxygen binding affinity [42]. However, the significant downregulation of E91D observed in this study indicates reduced oxygen affinity of HBB in lung cancer tissues. Hypoxia is a well-established hallmark of multiple human malignancies, particularly lung cancers [43, 44], and this substitution may directly contribute to the formation and maintenance of the hypoxic tumor microenvironment.

In summary, systematic mapping of AA substitutomes in these key exosomal proteins uncovers novel cancer-specific substitution signatures, providing mechanistic insights into tumor progression and identifying promising candidate biomarkers and therapeutic targets for further preclinical validation.

#### 2.2.2 AA substitutomes in actin filament binding proteins across cancer cohorts

Actin is the major component of the cytoskeletal protein in most cells. The polymerization of actin forms the actin filament (microfilament), which is seven nanometers in diameter and several micrometers in length [45]. In cells, the microfilaments are highly organized and form structures of semisolid gels. Various actin-binding proteins rule the assembly and disassembly of actin filaments and are, therefore, important to the actin skeleton. It has been widely found that actin-binding proteins have associations with cancer progression and pave the way to possible therapeutic methods [46, 47, 48].

In our study, GO enrichment analysis shows that, based on the up-regulated UPSPs in the five cancer cohorts, the proteins are most abundantly enriched in the functions of protein binding and RNA binding. Among the five cancers, four types of cancers, except for the ccRCC, are functionally enriched in actin filament binding. In the functional enrichment results with more than five enriched genes, 13 out of 15 in GBM, 3 out of 6 in HNSCC, 6 out of 7 in LUAD, 5 out of 7 in LSCC, and 1 out of 1 in ccRCC enriched functions are shown to be related to the binding function (Figure 4A). The AA substitutions can alter the primary structure of involved proteins, thereby changing the protein fold and intra-molecular bonds of the AA chain. These changes work together to influence the binding sites and functions [49]. This observation deepens the motivation of our AA substitutions investigation. The GO molecular function enrichment results based on the down-regulated UPSPs are shown in Supplementary Figure S12.

**Figure 4:**
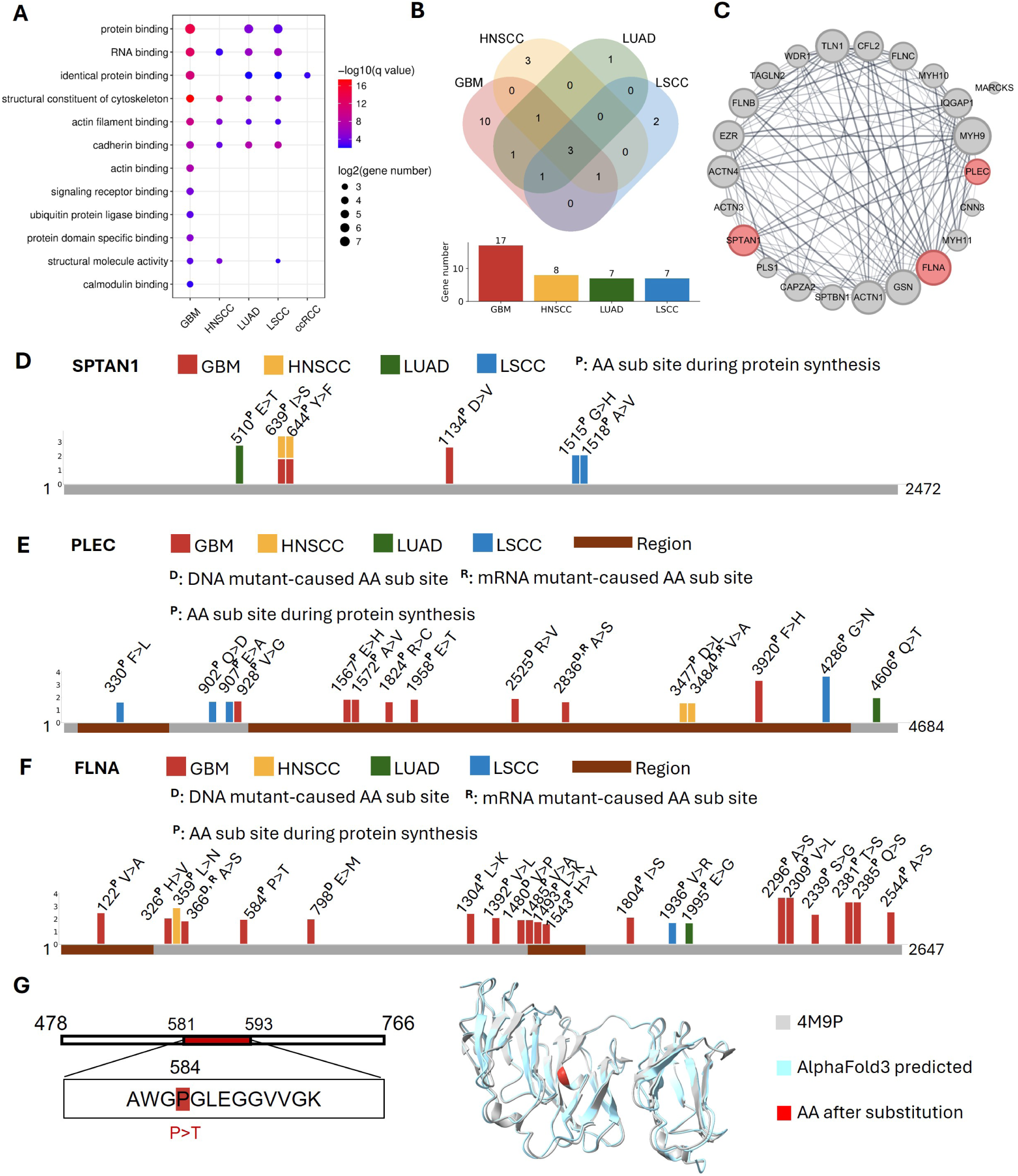
AA substitutomes in actin filament binding proteins across cancer cohorts. (A) Bubble plot showing the molecular function terms of GO enrichment based on the upregulations in the five cancer cohorts. The size of the bubble represents the base two logarithm of the enriched gene number, and the color of the bubble shows the minus base 10 logarithm of the q-value. (B) Venn plot (top) and bar plot (bottom) demonstrating the number of genes related to actin filament binding in four cohorts of cancers. The cohorts are GBM (dark red), HNSCC (yellow), LUAD (green), and LSCC (blue). (C) Diagram representing a network of 23 proteins corresponding to actin filament binding that are significantly up-regulated. The size of the node represents the degree of the protein. (D) Cumulative AA substitutions map of PLEC across different cancer cohorts. The AA substitutions above the protein sequence are up-regulated (Supplementary File 4). Barplot showing the base 2 logarithm ratio of cancer regulation. ^P^: AA substitution site during protein synthesis. The cohorts are GBM (dark red), HNSCC (yellow), LUAD (green), and LSCC (blue). (E) Cumulative AA substitutions map of PLEC across different cancer cohorts. The AA substitutions above the protein sequence are up-regulated (Supplementary File 4). Barplot showing the base 2 logarithm ratio of cancer regulation. Brown segment: binding and interaction region. ^D^: DNA mutant-caused AA substitution site. ^R^: mRNA mutant-caused AA substitution site. ^P^: AA substitution site during protein synthesis. The cohorts are GBM (dark red), HNSCC (yellow), LUAD (green), and LSCC (blue). (F) Cumulative AA substitutions map of FLNA across different cancer cohorts. The AA substitutions above the protein sequence are up-regulated (Supplementary File 4). Barplot showing the base 2 logarithm ratio of cancer regulation. Brown segment: binding and interaction region. ^D^: DNA mutant-caused AA substitution site. ^R^: mRNA mutant-caused AA substitution site. ^P^: AA substitution site during protein synthesis. The cohorts are GBM (dark red), HNSCC (yellow), LUAD (green), and LSCC (blue). (G) Structure comparison between the X-ray diffraction structure of FLNA Ig-like domains 3–5 (PDB: 4M9P) and structure prediction with AA substitution of P584T from the GBM cohort. Left: the site of proline to threonine (P584T) substitution in the protein sequence. Right: the structure of 4M9P (gray) and AlphaFold3-predicted structure after P584T substitution (light blue). The AAs after substitution (red).

The number of actin filament binding-related genes of each cancer cohort and the gene overlap is displayed in Figure 4B, where the colors correspond to four of the cancer types, excepting ccRCC. Among the four cancers, 23 proteins are actin filament binding-related and GBM contributes 17 proteins. To demonstrate the PPI relationships between the microfilament binding proteins that are expressed and identified in the cancer cohorts, we show the 23 proteins in Figure 4C, where the size reflects the degree of each protein.

Spectrins represent a group of filament cytoskeletal proteins, which serve as crucial scaffold proteins, playing a vital role in supporting the plasma membrane and arranging internal organelles. Spectrins consist of *α* and *β* chains that come together to create tetramers arranged in a head-to-head style. SPTAN1 encodes the *α* chain of spectrin and is exclusively found in non-erythrocytic cells [50]. With its diverse functions encompassing adhesion and migration, SPTAN1 has both positive and negative impacts on tumor development and progression, depending on the regulation situation of spectrin alpha chain (SPTAN1) [51]. In the AA substitutions analysis results of SPTAN1, we find four AA substitutions in GBM, two in HNSCC, one in LUAD, and two in LSCC. Additionally, two common AA substitutions, I639S substitution and Y644F substitution, are identified from GBM and HNSCC (Figure 4D, Supplementary File 2). The UPSPs with common AA substitutions show a 3.56-fold and 3.07-fold increase in abundance in GBM and HNSCC, respectively.

Plectin (PLEC), a long-sequence protein present in almost all mammalian cells, serves as a crucial connection point among the primary components of the cytoskeleton proteins, including actin microfilaments [52]. In PLEC, more AA substitutions are identified from the four cancer cohorts. We point out eight up-regulated AA substitutions in GBM, two in HNSCC, one in LUAD, and four in LSCC. Among these AA substitutions sites, sites 2525, 2836, and 3484 are three sites recorded to have AA substitutions caused by DNA or mRNA mutation. R2525C is found to be caused by DNA mutation, while A2836T and V3484M are caused by both DNA and mRNA mutation, according to OncoDB [37] (Supplementary File 4). Two binding and interaction regions in the PLEC sequence cover most of the identified AA substitutions (Figure 4E). The first region ranges from 175–400, which is an actin-binding region [53], and in the LSCC cohort we find an F330L substitution, which replaces an aromatic AA with a non-aromatic AA, in this region. The second region includes the sequence of 964–4574, which covers about 77% of PLEC’s length. This region has been previously discovered to interact with IF proteins [54]. It includes seven AA substitutions in GBM, two AA substitutions in HNSCC, and one in LSCC, and all of the covered AA substitutions are upregulated substitutions. The interaction between PLEC and IF proteins might be influenced by the AA substitutions crowding in this region.

Filamin-A (FLNA) is closely involved with the proliferation and invasion of GBM [55, 56], and has been studied together with the mechanistic target of rapamycin complex two activity as a signaling pathway in the motility of GBM cells [57]. Meanwhile, FLNA is an essential protein in linking actin filaments to integrate molecules within cell membranes, crucially contributing to maintaining cell shape and signaling processes [58]. Probing the AA substitutions in GBM and GBM-regulated UPSPs potentially creates a new angle to uncover the mechanism of migration and invasion of GBM cells. The number of up-regulated AA substitutions sites of FLNA in the GBM cohort is 19. Among these substitution sites G776V, E798K, and V1480M have been recorded as DNA mutation-caused AA substitutions, while A366T and G776S have been recorded to be caused by both DNA mutation and mRNA mutation (Supplementary File 4). The rest of the identified AA substitutions should take place after protein translation. Interestingly, substitution sites 1804 and 2544 are identified to have two possible AA substitutions in GBM (Figure 4F, Supplementary File 2). The number of substitutions for the other three cohorts is just one. Some AA substitutions are located in the regions of binding and interaction. In the actin-binding region from 2 to 274, there is a V122A substitution. In the region of interaction with furin from sequence 1490 to 1607, two AA substitutions (L1493K and H1543Y) are identified.

The FLNA dimer structure has 24 immunoglobulin-like (Ig) domains, among which domain 4 is stabilized by interaction with domain 5 and the interface between domain 4 and domain 5 is governed by several hydrophobic *β*-sheet interactions [59]. Additionally, two AA substitutions in domains 4 and 5, P637E and V711D, respectively, have been previously reported to cause familial cardiac valvular dystrophy because these two substitutions likely change the hydrophobic core and destabilize the individual *β*-sheet domain formulation [60].

One of the AA substitutions identified in FLNA has been validated by a previous study, highlighting significant biological implications. Our analysis reveals a P584T substitution in FLNA domain 4, which is a hydrophobic-to-hydrophilic substitution with a four-fold up-regulation (Supplementary File 2). Although this residue does not reside at domain interfaces, its side chain orientation toward the *β*-sheet hydrophobic core suggests that P>T substitution may substantially perturb core packing, potentially destabilizing the *β*-sheet domain architecture. This conformational disruption likely disorders the FLNA-integrin interaction interface critical for *β*-sheet stabilization. Such structural alterations may underlie the observed enhancement of mutant FLNA’s proliferative and invasive functions in GBM, since the integrin activation is essential for cellular propulsion following FLNA release [61]. To evaluate these structural changes, we employ AlphaFold3 [62] to model the P584T variant and compared it with the FLNA domains 3-5 X-ray diffraction structure [59] (Figure 4G), providing mechanistic insights into the mutation’s functional consequences. These findings provide mechanistic insights into how specific AA substitutomes may contribute to cancer progression through structural perturbation of key cytoskeletal proteins.

#### 2.2.3 AA substitutomes in glycolysis and gluconeogenesis proteins across cancer cohorts

Glycolysis and gluconeogenesis are two reciprocally regulated pathways. Glucose is broken down to pyruvate in the glycolysis pathway and a little adenosine triphosphate (ATP) is made in this process. Gluconeogenesis is similar to the reverse process of glycolysis, producing glucose de novo from precursors, including pyruvate or alanine. The pathway of gluconeogenesis is closed when glycolysis is stimulated and vice versa [63]. It has been revealed that many cancer cells take glycolysis as an essential source of energy, because of metabolic and mitochondrial imperfection. Additionally, the speed of ATP production by glycolysis is faster than mitochondrial oxidative phosphorylation. Therefore, it is more suitable for the rapid energy demands of cancer cell proliferation [64]. At the same time, the simultaneous activation or suppression of glycolysis and gluconeogenesis could result in metabolic stress and lead to metabolic interference [65]. Therefore, the study of these pathways could provide therapeutic approaches to cancers due to their pivotal role in cancer cell proliferation and viability. KEGG pathways enrichment [30] among the five cancer cohorts shows that the AA substitutions-containing proteins are enriched in the pathways of glycolysis and gluconeogenesis. For the up-regulations, four of the cancer cohorts, excepting HNSCC, are enriched in these pathways (Figure 5A, indicated by the arrow), while in the down-regulations, the results of ccRCC are enriched (Figure 5B, indicated by the arrow).

**Figure 5:**
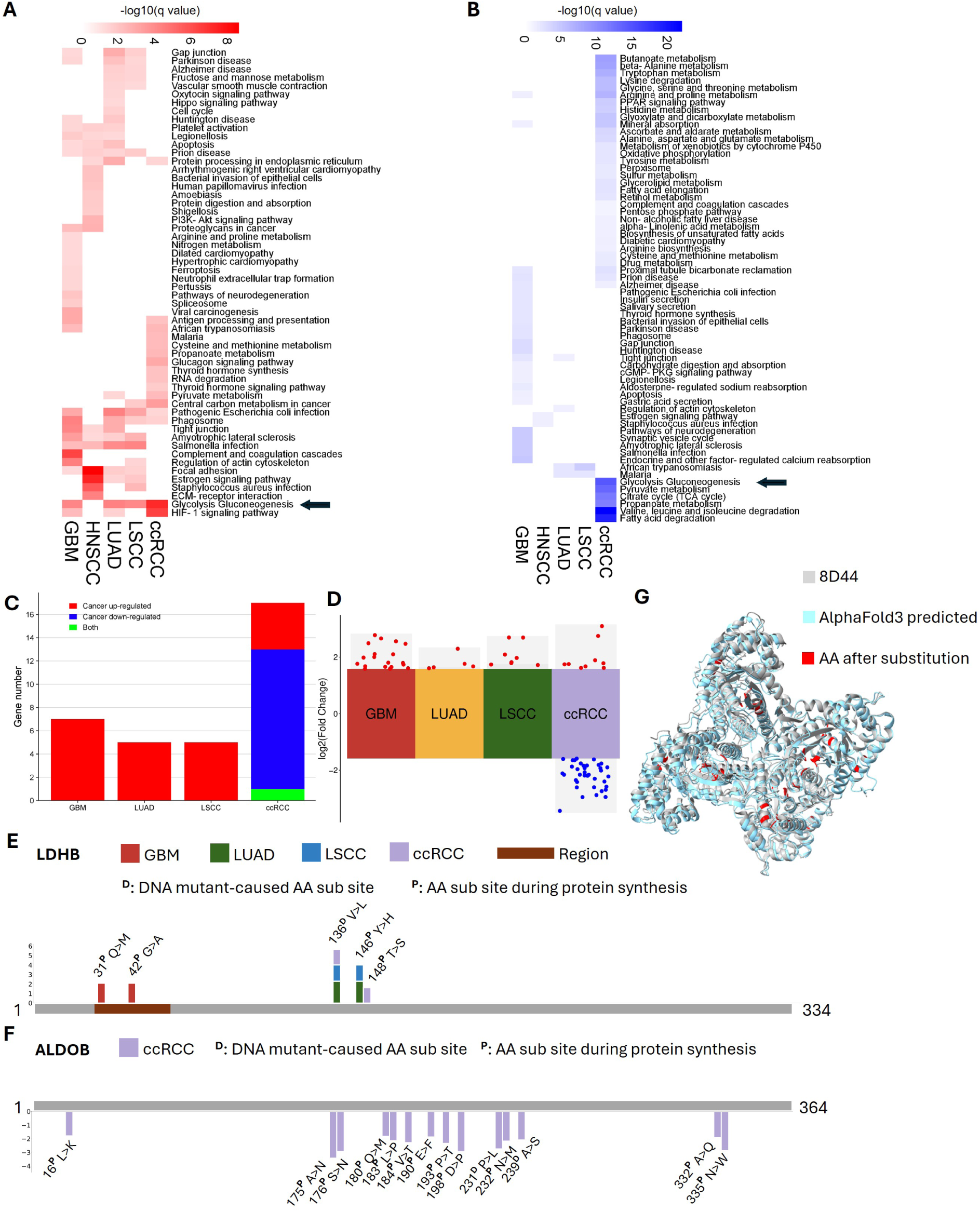
AA substitutomes in glycolysis and gluconeogenesis proteins across cancer cohorts. (A) Heatmaps showing KEGG enrichment results of up-regulations among five cancer cohorts. The color represents the minus base 10 logarithm of the q-value; the darker the red, the smaller the q-value. (B) Heatmaps showing KEGG enrichment results of down-regulations among five cancer cohorts. The color represents the minus base 10 logarithm of the q-value; the darker the blue, the smaller the q-value. (C) Bar plot showing the number of genes related to glycolysis and gluconeogenesis among four types of cancers. The up-regulated-only genes are denoted by the red segment, the down-regulatedonly genes are denoted by the blue segment, and the both upand down-regulated genes are denoted by the green segment. (D) Volcano plot presenting the gene expression of glycolysis and gluconeogenesis corresponding UPSPs. The cohorts are GBM (dark red), LUAD (green), LSCC (blue), and ccRCC (purple). (E) Cumulative AA substitutions map of LDHB in four types of cancers. The AA substitutions above the protein sequence are up-regulated (Supplementary File 4). Barplot showing the base 2 logarithm ratio of cancer regulation. Brown segment: binding and interaction region. ^D^: DNA mutant-caused AA substitution site. ^P^: AA substitution site during protein synthesis. The cohorts are GBM (dark red), LUAD (green), LSCC (blue), and ccRCC (purple). (F) Cumulative AA substitutions map of ALDOB in ccRCC. The AA substitutions below the protein sequence are down-regulated (Supplementary File 4). Barplot showing the base 2 logarithm ratio of cancer regulation. ^D^: DNA mutant-caused AA substitution site. ^P^: AA substitution site during protein synthesis. The cohort is ccRCC (purple). (G) The structure comparison between the cryo-electron microscopy structure of human renal ALDOB (PDB: 8D44) and structure prediction with cumulative AA substitutions from ccRCC. Gray: the structure of 8D44. Light blue: AlphaFold3-predicted structure with cumulative AA substitutions. Red: The AAs after substitution.

The numbers of glycolysis and gluconeogenesis pathways-related genes are displayed in Figure 5C. Among the four cohorts, GBM has seven up-regulated genes, which is more than the other cancer types, while ccRCC has 12 down-regulated genes and one bidirectionally regulated gene, and is the only cancer that has down-regulated AA substitutions. The regulation situation of the significantly regulated UPSPs of the four cohorts is demonstrated in Figure 5D. We observe that the L-lactate dehydrogenase B chain (LDHB) is mutually expressed by the four cancers (GBM, LUAD, LSCC, and ccRCC) in our analysis results. In the glycolysis pathway, LDHB holds a vital position because it is a crucial glycolytic enzyme responsible for catalyzing the conversion of pyruvate to lactate and the backward conversion [66]. And since the enzymatic activity and direction of LDHB dictate the pace of glycolysis, LDHB is under active regulation in cancer cells [67]. While the critical involvement of LDHB in the advancement of different cancers has been increasingly documented [68, 69], the AA substitutomics analysis of LDHB across different cancers remains inadequate. Besides, among the down-regulated proteins, fructose-bisphosphate aldolase B (ALDOB) is the one that is modified by the largest number of AA substitutions. ALDOB is a type of glycolytic enzyme that facilitates the transformation of fructose 1,6-diphosphate into glyceraldehyde 3-phosphate and dihydroxyacetone phosphate within the glycolysis metabolic pathway [70]. As a possible prediagnosis indicator for ccRCC patients, the down-regulated expression of ALDOB has been strongly linked to clinicopathological characteristics of ccRCC patients [71]. The literature validates the findings in this study of down-regulated AA substitutions in ALDOB from the ccRCC cohort.

We continue to analyze the profile of the AA substitutions for the AA substitutions-modified, glycolysis and gluconeogenesis pathways-enriched proteins LDHB and ALDOB. Among the identified AA substitutions and substitution sites in LDHB and ALDOB, some AA substitutions sites have been found to have DNA mutation-caused AA substitutions [37]. Specifically, in LDHB, the DNA mutation-caused AA substitution is V136M, and in ALDOB, the DNA mutation-caused AA substitutions are P231R and A239S (Supplementary File 4). The up-regulated AA substitutions identified in the four cohorts are annotated in Figure 5E, showing two AA substitutions for each cancer cohort (Supplementary File 2). The two AA substitutions of LDHB from the GBM cohort reside in the binding region of the oxidized form of nicotinamide adenine dinucleotide (NAD^+^), whose sequence ranges from 31–53. LDHB can catalyze the reversible transformation of lactate and NAD^+^ into pyruvate and NADH (H for hydrogen) [72]. The two AA substitutions could interfere with the conversion process and the glycolysis and gluconeogenesis pathways and affect the energy generation in cancer cells.

ALDOB emerges as another protein with frequent substitutions, and one of the identified AA substitution sites in ALDOB have been validated by other research studies. Notably, we report the novel identification of an A175N substitution at a well-documented mutational hotspot [73, 74, 75] (Figure 5F). Position 175 contains an evolutionarily conserved alanine across all aldolase isozymes, located within an *α*-helical region proximal to substrate-binding residues according to crystallographic studies [76]. Substituting alanine at this position alters the charge of the AA, which may disrupt the spatial configuration of neighboring residues on the *β*-strand of the ALDOB substrate-binding pocket responsible for ATP generation. Moreover, the A>N substitution modifies the intra-chain hydrogen bond between sites 175 and 179 while also changing the hydrophilicity of residue 175 [77]. These alterations potentially impact molecular stability and consequently influence the glycolytic and gluconeogenic pathways, ultimately impairing energy production in cancer cells. All identified AA substitutions in ALDOB are exclusively found within the renal cancer cohort and exhibit a consistent down-regulation (Supplementary File 2). This finding aligns with previous research, which links the down-regulation of ALDOB to pathological characteristics observed in ccRCC patients [71].

Recently, the experimental structure of human kidney ALDOB was proposed (PDB: 8D44). To exhibit the influence of the AA substitutions in this study on protein structure, we present a structural comparison between the experimental structure and the AlphaFold3-predicted structure with cumulative AA substitutions (Figure 5G). Molecular analysis of these newly discovered AA substitutions and function analysis of the AA substitutions-modified ALDOB remain to be performed. These findings provide novel insights into how AA substitutomes may contribute to metabolic reprogramming in cancer. Further functional characterization of these substitutions will be essential to fully elucidate their roles in cancer metabolism.

### 2.3 AA substitutomic explains the drug escape mechanism of cancer

AA substitutions in cancer cells represent a crucial mechanism contributing to drug escape [78], as mutations in essential metabolic enzymes or transporters modify protein structure and function, thereby facilitating escape from therapeutic agents [79]. A wealth of research has underscored the importance of AA substitutions in the context of cancer drug escape. For example, mutations in enzymes such as glutaminase and asparagine synthetase can enhance their stability or substrate affinity, enabling cancer cells to sustain vital biosynthetic and redox balance pathways despite targeted drug interventions [80, 81]. These substitutions often induce structural alterations in active sites or allosteric regulatory domains, which can disrupt drug binding while maintaining enzymatic activity. In the case of cisplatin-escape cancers, for instance, the upregulation of glutamine metabolism supports nucleotide synthesis and glutathione production, thereby alleviating drug-induced oxidative stress [80]. Likewise, mutations in AA transporters may increase their affinity for substrates such as cystine or leucine, promoting antioxidant defense mechanisms and mTOR-driven proliferation even in the presence of therapeutic inhibition [82]. Such substitutions frequently arise under conditions of drug pressure, illustrating how cancer cells leverage AA metabolism to circumvent therapeutic strategies.

In this study, the AA substitutomic analysis pipeline successfully identifies four AA substitutions in low-density lipoprotein receptor-related protein 2 (LRP2) from the ccRCC cohort (Supplementary File 2, Figure 6A, Supplementary Figure S13). LRP2, also referred to as megalin, is recognized as the largest known endocytic membrane receptor, playing a crucial role in the endocytosis of various ligands in specialized epithelial tissues, such as the proximal tubules of the kidney. Previous reports have indicated that low expression levels of LRP2 correlate with poor patient outcomes in ccRCC [83], a finding that aligns with our observations in this study. Furthermore, LRP2’s significance extends to targeted therapies, particularly concerning vitamin B12 uptake and its potential use as a vector for drug delivery [84]. Given the extensive length of LRP2 and the fact that the latest structural model (PDB: 9CWM) does not provide a continuous representation, we opted to enhance visualization by presenting the structural comparisons within segments 3501–3700 (Figure 6B left) and 4201–4300 (Figure 6B right), which encompass the identified AA substitutions. The comparison between the experimental structure and the AlphaFold3-predicted model, incorporating cumulative AA substitutions, suggests that the identified substitutions may have a tentative impact on the protein structure of LRP2. Although specific therapeutic antibodies targeting LRP2 are not yet widely available, there is ongoing investigation into antibodies against LRP2 for their potential applications in cancer diagnosis and therapy. For example, the presence of LRP2 on epithelial cell surfaces makes it an appealing target for therapeutic antibodies [83]. The utilization of AA substitutomics enables a detailed investigation of LRP2 AA substitutions in cancer, ultimately aiding in the formulation of innovative therapeutic strategies aimed at targeting LRP2 while circumventing the associated drug escape mechanisms.

**Figure 6:**
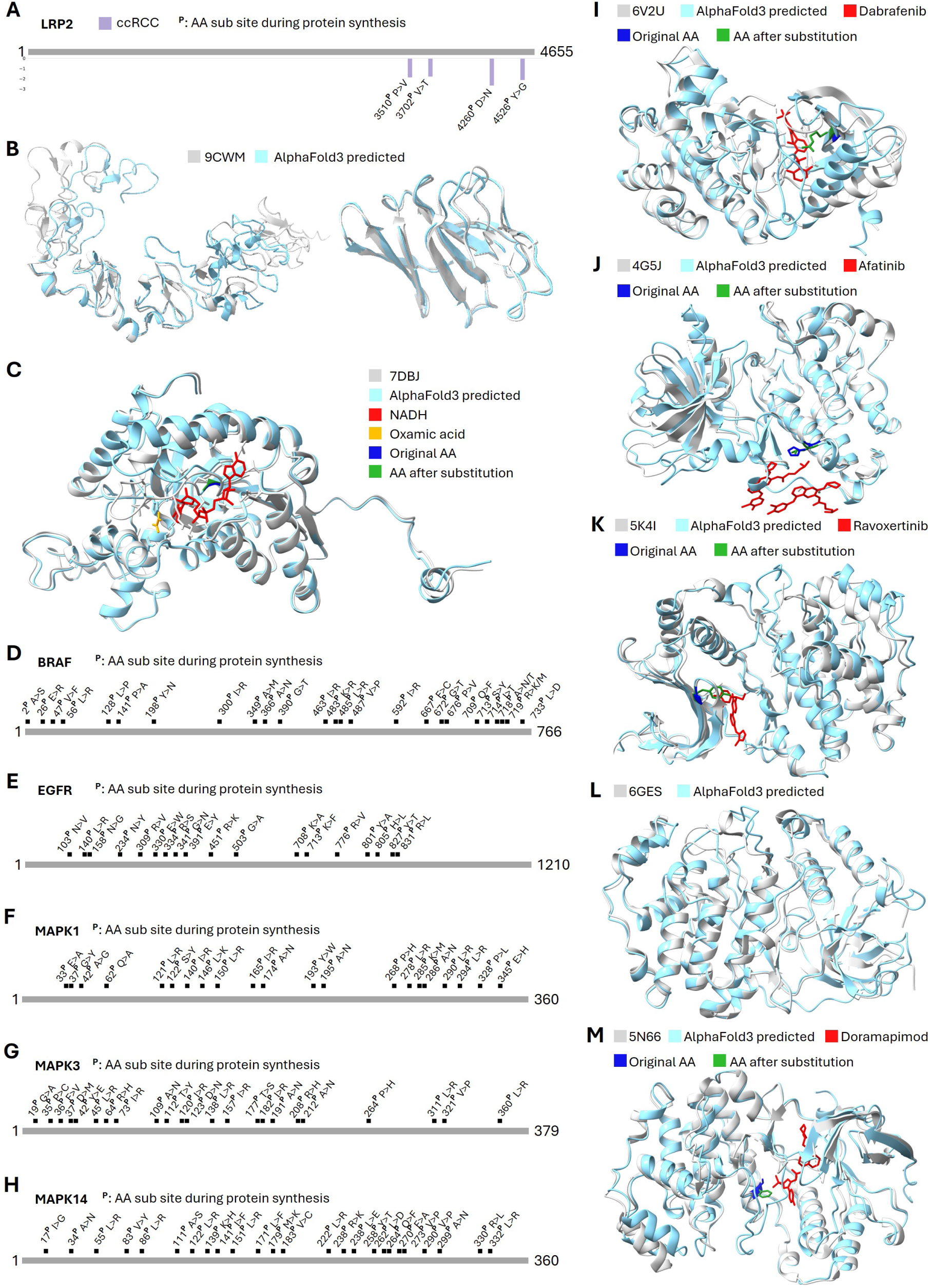
AA substitutomic explains the drug escape mechanism of cancer. (A) Cumulative AA substitutions map of LRP2 in the ccRCC cohort (purple). The AA substitutions below the protein sequence are down-regulated. Barplot showing the base 2 logarithm ratio of cancer regulation (Supplementary File 2). ^P^: AA substitution site during protein synthesis. (B) The structure comparison between the cryo-electron microscopy structure of human LRP2 (PDB: 9CWM) and the structure prediction of LRP2 with cumulative AA substitutions in the segment of 3501–3700 (left) and 4201–4300 (right). Gray: the structure of 9CWM. Light blue: AlphaFold3-predicted structure. (C) The structure comparison between the X-ray diffraction structure of human LDHB (PDB: 7DBJ) and the structure prediction of LDHB with cumulative AA substitutions. Gray: the structure of 7DBJ. Light blue: AlphaFold3-predicted structure. Red: oxamic acid. Orange: NADH. Deep blue: the original AA. Green: the AA after substitution. (D) Cumulative AA substitutions map of BRAF in the BRAF enriched dataset (Supplementary File 5). ^P^: AA substitution site during protein synthesis. (E) Cumulative AA substitutions map of EGFR in the EGFR enriched dataset (Supplementary File 5). ^P^: AA substitution site during protein synthesis. (F) Cumulative AA substitutions map of MAPK1 in the MAPK1 enriched dataset (Supplementary File 5). ^P^: AA substitution site during protein synthesis. (G) Cumulative AA substitutions map of MAPK3 in the MAPK3 enriched dataset (Supplementary File 5). ^P^: AA substitution site during protein synthesis. (H) Cumulative AA substitutions map of MAPK14 in the MAPK14 enriched dataset (Supplementary File 5). ^P^: AA substitution site during protein synthesis. (I) The structure comparison between the X-ray diffraction structure of human BRAF (PDB: 6V2U) and the structure prediction of BRAF with cumulative AA substitutions. Gray: the structure of 6V2U. Light blue: AlphaFold3-predicted structure. Red: Dabrafenib. Deep blue: the original AA. Green: the AA after substitution. (J) The structure comparison between the X-ray diffraction structure of human EGFR (PDB: 4G5J) and the structure prediction of EGFR with cumulative AA substitutions. Gray: the structure of 4G5J. Light blue: AlphaFold3-predicted structure. Red: Afatinib. Deep blue: the original AA. Green: the AA after substitution. (K) The structure comparison between the X-ray diffraction structure of human MAPK1 (PDB: 5K4I) and the structure prediction of MAPK1 with cumulative AA substitutions. Gray: the structure of 5K4I. Light blue: AlphaFold3-predicted structure. Red: Ravoxertinib. Deep blue: the original AA. Green: the AA after substitution. (L) The structure comparison between the X-ray diffraction structure of human MAPK3 (PDB: 6GES) and the structure prediction of MAPK3 with cumulative AA substitutions. Gray: the structure of 6GES. Light blue: AlphaFold3-predicted structure. (M) The structure comparison between the X-ray diffraction structure of human MAPK14 (PDB: 5N66) and the structure prediction of MAPK14 with cumulative AA substitutions. Gray: the structure of 5N66. Light blue: AlphaFold3-predicted structure. Red: Doramapimod. Deep blue: the original AA. Green: the AA after substitution.

In addition to LRP2, this study identifies LDHB as a protein potentially involved in drug escape mechanisms in cancer. During aerobic glycolysis, the conversion of pyruvate to lactate is facilitated by cytosolic LDH enzymes, utilizing NADH as a cofactor, which is subsequently converted to NAD^+^ [72]. LDHB has been linked to aggressive phenotypes in lung cancer [68] and plays a significant role in lysosomal activity and autophagy within cancer cells [69]. Given that many cancer cells upregulate autophagy to support metabolism, tumorigenesis, and resistance to therapies [85], targeting LDHB inhibition presents a promising approach for the prevention and treatment of various cancers. In this research, we identified five AA substitutions across four cohorts (GBM, LUAD, LSCC, and ccRCC) that may impact the structure of LDHB (Figure 6C). The X-ray diffraction structure of the quaternary complex of LDHB with NADH and oxamic acid elucidates the spatial relationship and binding interface between LDHB and these compounds [86]. Oxamic acid has been reported to exhibit inhibitory effects on LDH [87]. The NADH-oxamic acid complex forms hydrogen bonds with LDHB, with asparagine at site 138 being one of the binding AAs. The AA substitutomic analysis pipeline has revealed a V136L substitution located near this binding asparagine, which could potentially affect the binding affinity of LDHB. While NADH and oxamic acid are not classified as cancer drugs, NADH metabolism is known to be involved in the growth and survival of cancer cells [88], and oxamic acid demonstrates potential anti-cancer properties, particularly against aggressive tumors such as brain cancers [89].

Nowadays, numerous proteins have been identified as drug targets for cancer therapeutics, including serine/threonine-protein kinase B-raf (BRAF), epidermal growth factor receptor (EGFR), and Mitogen-activated protein kinase (MAPK) proteins, etc [90]. Identifying AA substitutions in these drug target proteins is crucial for understanding the role of these substitutions in drug escape mechanisms. However, across the five cohorts analyzed, we observed that such cancer drug target proteins are not abundant, especially when compared to the eight proteins discussed in section 2.2 (Supplementary Figure S13). Consequently, we collected enriched datasets of cancer drug target proteins to further explore this aspect [91, 92]. The dataset is collected from ProteomeXchange, utilizing the following data identifiers: PXD000788 [91], PXD016465, PXD016505, and PXD016463 [92]. The search results are available on Zenodo, with record number 17669590. Figure 6D-H (Supplementary File 5) presents the cumulative AA substitution maps for BRAF, EGFR, MAPK1, MAPK3, and MAPK14, respectively. Additionally, Figure 6I-M illustrates the structural comparisons between the original protein structures and the AlphaFold3 predicted structures for these drug targets, highlighting the impact of the substituted AAs on drug binding interactions. Therefore, we focus on the structural comparison between the original and the AlphaFold predicted structure, showcasing the cumulative AA substitutions identified. Besides, to quantify the impact of AA substitutions on drug-target interactions, we utilized CB-Dock2 [93] to simulate docking between the drugs and their respective protein targets before and after AA substitution modification. Detailed instructions for using CB-Dock2 are provided in the Methods section. CB-Dock2 employs the Vina score to assess the binding affinity between each drug molecule and its targeted protein, with lower scores indicating stronger predicted binding affinities.

The BRAF protein is a key regulator in the MAPK/ERK signaling pathway, which transmits mitogenic signals from cell surface receptors to the nucleus. When activated, BRAF phosphorylates and activates MEK, leading to a kinase cascade that regulates essential cellular functions such as proliferation, differentiation, and survival [94]. Dabrafenib is a selective ATP-competitive inhibitor that targets the kinase activity of BRAF proteins with the V600E mutation, which continuously activates the MAPK/ERK pathway and drives unchecked cellular proliferation and survival [95]. In this study, we identify a K483R substitution that may significantly reduce binding affinity by disrupting a key electrostatic interaction in the dabrafenib-BRAF interface (Figure 6I). Normally, lysine 483 forms an essential salt bridge with the negatively charged sulfonamide group of dabrafenib. The substitution to arginine, while still positively charged, introduces steric and geometric constraints that hinder this interaction. The bulkier guanidinium group of arginine requires different spatial orientation for effective hydrogen bonding compared to lysine’s primary amine [96]. And average Vina scores changed from -11.2 to -10.6 for the Dabrafenib-BRAF pair (Supplementary Figure S14). This disruption likely compromises the drug’s binding affinity, leading to a decreased inhibitory potency.

EGFR is a transmembrane receptor tyrosine kinase that plays a vital role in regulating cellular proliferation, survival, and differentiation upon activation. Its significance is highlighted by the frequent oncogenic activation due to gain-of-function mutations in the kinase domain, which promote uncontrolled growth in cancers such as LUAD and LSCC [97]. Afatinib is a second-generation, irreversible tyrosine kinase inhibitor that covalently binds to a conserved cysteine residue at site 797 in the ATP-binding pocket of EGFR, providing strong and sustained inhibition of oncogenic signaling [98]. The identified H805L mutation (Figure 6J) occurs within the kinase domain and is believed to reduce binding affinity by disrupting crucial molecular interactions necessary for maintaining the drug-protein complex. Histidine 805 likely contributes to a network of hydrogen bonds and aids in the precise orientation of the binding pocket [99]. Substituting this histidine with the hydrophobic, sterically distinct leucine alters local electrostatics and van der Waals contacts, possibly inducing a conformational shift that compromises the positioning of afatinib for its covalent interaction with the conserved cysteine. And the average Vina scores changed from -8.6 to -7.9 for the Afatinib-EGFR pair (Supplementary Figure S14). This disruption may reduce the drug’s inhibitory efficacy and facilitate escape from targeted therapy.

MAPK1 serves as a terminal serine/threonine kinase in the MAPK/ERK signaling cascade, transmitting mitogenic and survival signals from upstream regulators to a wide range of cytosolic and nuclear substrates. This function is essential for regulating cell proliferation, differentiation, and survival, with its dysregulation being a key factor in oncogenesis [100]. Ravoxertinib is a potent and selective ATP-competitive inhibitor designed to bind to the active conformation of MAPK1, occupying the kinase’s catalytic cleft to inhibit phosphotransferase activity and counteract the protumorigenic effects of the hyperactivated pathway [101]. However, the identified G37Y mutation (Figure 6K) directly undermines this therapeutic approach. Glycine 37 is a highly conserved residue in the phosphate-binding loop, and its replacement with the bulky, aromatic side chain of tyrosine introduces significant steric hindrance, potentially altering the structure of the ATP-binding pocket [102]. This steric clash obstructs the optimal docking of ravoxertinib, thereby diminishing its binding affinity and allowing continued kinase activity. And the average Vina scores changed from -9.0 to -8.4 for the Ravoxertinib-MAPK1 pair (Supplementary Figure S14). From a drug escape standpoint, this mutation offers a direct mechanism, as it selectively impedes inhibitor binding while likely maintaining the kinase’s capacity to bind its natural ATP substrate and downstream effectors, thus facilitating tumor cell survival and proliferation even in the presence of the drug.

MAPK14 is a stress-activated serine/threonine kinase that plays a crucial role in mediating inflammatory responses, regulating the cell cycle, and promoting apoptosis. Its dysregulated activity has been linked to the pathogenesis of autoimmune diseases and multiple cancers, highlighting its significance as a therapeutic target [103]. Doramapimod is a potent and selective ATP-competitive inhibitor of MAPK14 that employs a unique mechanism by not only occupying the ATP-binding pocket but also stabilizing a specific DFG-out conformation, locking the kinase in an inactive state [104].In this study, we identify the I141F substitution (Figure 6M), which directly undermines this mechanism. Isoleucine 141 is a key hydrophobic residue within the allosteric pocket adjacent to the ATP-binding site that accommodates the large, rigid scaffold of doramapimod. The substitution with the bulkier and conformationally constrained phenylalanine introduces significant steric hindrance and alters the local hydrophobic environment [105]. This disruption impedes the essential van der Waals interactions required for high-affinity drug binding and hampers the stable induction of the DFG-out state. And the average Vina scores changed from -10.7 to -10.2 for the Doramapimod-MAPK14 pair (Supplementary Figure S14). Consequently, this mutation exemplifies a direct allosteric resistance mechanism, where a single AA substitution in the inhibitor’s binding region markedly decreases binding affinity without compromising the kinase’s intrinsic catalytic competence for ATP.

The exploration of AA substitutions at the binding interfaces of oncogenic kinases and their targeted inhibitors offers valuable insights into the mechanisms of drug escape. In each instance, whether through steric hindrance, disruption of crucial electrostatic or hydrophobic interactions, or alteration of allosteric conformational dynamics, these mutations significantly reduce binding affinity, enabling oncogenic kinases to evade pharmacological inhibition. The findings across various kinase-inhibitor pairs underscore the drug-binding site as a critical vulnerability where cancer cells can implement minimal AA changes to achieve substantial therapeutic escape. Therefore, the AA substitutomic analysis presented in this research could pave the way for investigating drug escape mechanisms and cancer progression, providing a structural framework for predicting escape mechanisms, informing the development of next-generation inhibitors, and personalizing combination therapies to circumvent these adaptive responses.

### 2.4 AA substitutomic reveals immune escape mechanism in cancer cell

Cancer immune escape is a pivotal hallmark of tumor progression, allowing malignant cells to avoid detection and elimination by the host’s immune system. A key aspect of this phenomenon is the altered presentation of tumor-associated antigens through major histocompatibility complex (MHC) molecules. Tumors employ intricate mechanisms to undermine antigen presentation, which include the selection of mutations that diminish peptide-MHC binding affinity [106]. A comprehensive understanding of these dynamics is crucial for the advancement of targeted immunotherapies designed to reinstate antitumor immunity [107].

AA substitutions in tumor-derived proteins constitute a crucial mechanism of immune escape by directly interfering with the antigen presentation pathway. These somatic mutations can significantly alter the peptide-MHC interaction by modifying critical anchor residues that are vital for stable binding within the MHC groove. Even a single substitution at these locations can substantially decrease the binding affinity of a peptide, thereby hindering its stable presentation on the tumor cell surface and rendering it undetectable to T cells [108]. Computational tools such as NetMHCpan have become essential for predicting peptide-MHC binding affinities and for identifying neoantigens—mutated peptides that are unique to tumors [109].

For those significantly AA substitution modified proteins from CPTAC cohorts, we find two proteins, PLEC and LDHB, that can bind with MHC proteins. Therefore, we compare the MHC binding affinities of the modified sequences with their corresponding wild-type sequences (Supplementary File 6). The outputs from NetMHCpan [109] reveal that, due to the AA substitutions, the number of MHC binders decreases from 90 to 86 for PLEC and from 8 to 7 for LDHB. This finding suggests a predicted reduction in binding affinity as indicated by NetMHCpan, which serves as a strong indicator that specific mutations may promote immune escape by hindering peptide presentation to T cells [109].

To further investigate the impact of AA substitutions on MHC proteins, we collected and analyzed the MHC protein-enriched datasets. The datasets are collected from ProteomeXchange with the identifiers of PXD003790 [110] and PXD000394 [111]. The search results are available on Zenodo, with record number 17669609. Figure 7A-D displays the cumulative identified AA substitution maps for human leukocyte antigen (HLA)-A, HLA-B, HLA-C, and HLA-DRB1, respectively (Supplementary File 7). Furthermore, Figure 7E-G presents structural comparisons between the original protein structures and the AlphaFold3 predicted models for these MHC proteins, emphasizing how the AA substitutions affect interactions with presented peptides. However, since AlphaFold3 was unable to accurately predict the structure of HLA-DRB1, we do not include its structural comparison in this paper.

**Figure 7:**
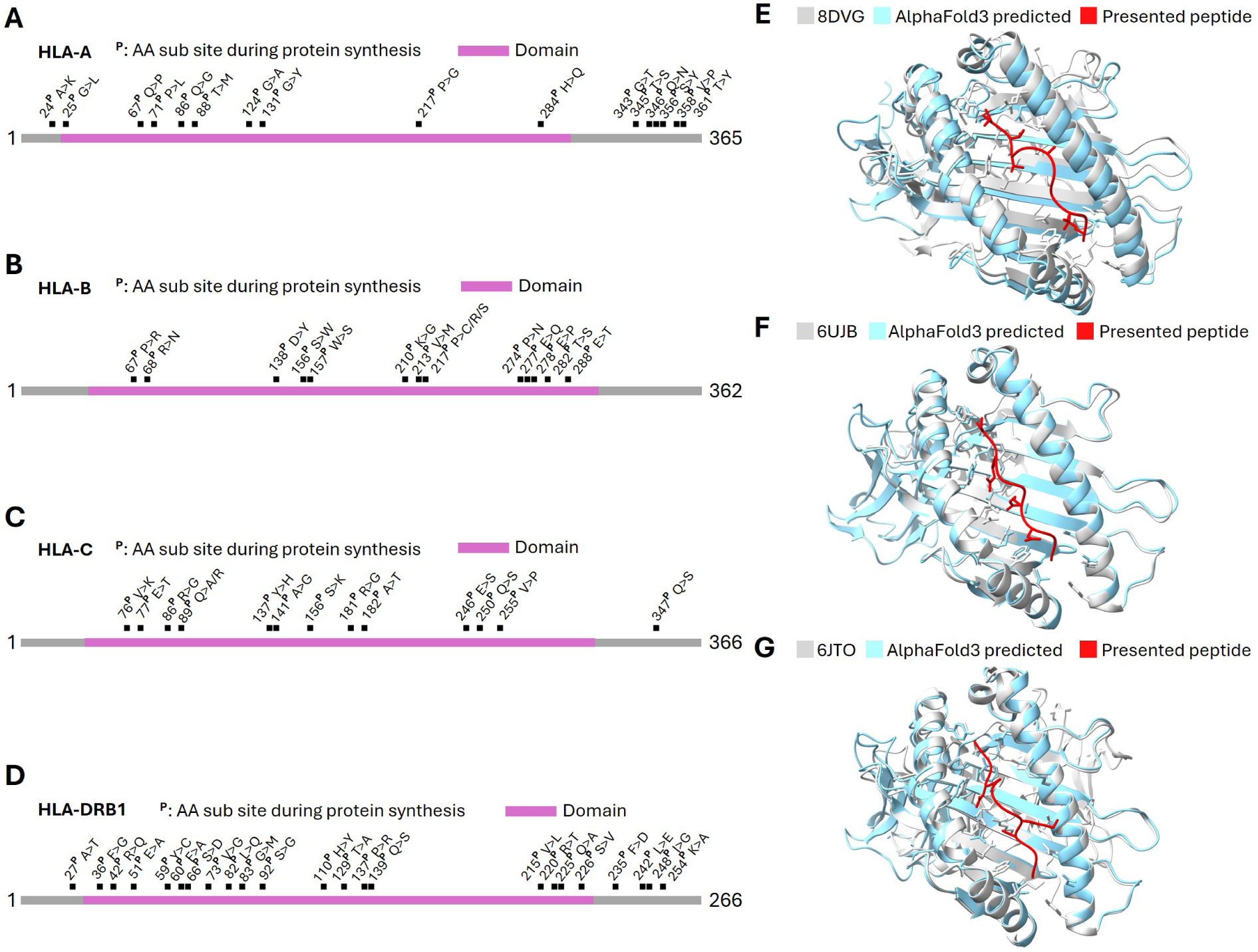
AA substitutomic explains the immune escape mechanism of cancer. (A) Cumulative AA substitutions map of HLA-A in the MHC protein enriched dataset (Supplementary File 7). ^P^: AA substitution site during protein synthesis. Pink segment: Alpha domains in the MHC class I protein. (B) Cumulative AA substitutions map of HLA-B in the MHC protein enriched dataset (Supplementary File 7). ^P^: AA substitution site during protein synthesis. Pink segment: Alpha domains in the MHC class I protein. (C) Cumulative AA substitutions map of HLA-C in the MHC protein enriched dataset (Supplementary File 7). ^P^: AA substitution site during protein synthesis. Pink segment: Alpha domains in the MHC class I protein. (D) Cumulative AA substitutions map of HLA-DRB1 in the MHC protein enriched dataset (Supplementary File 7). ^P^: AA substitution site during protein synthesis. Pink segment: Beta domains in the MHC class II protein. (E) The structure comparison between the X-ray diffraction structure of human HLA-A (PDB: 8DVG) and the structure prediction of HLA-A with cumulative AA substitutions. Gray: the structure of 8DVG. Light blue: AlphaFold3-predicted structure with cumulative AA substitutions. Red: presented peptide (VVVGAGGVGK). (F) The structure comparison between the electron microscopy structure of human HLA-B (PDB: 6UJB) and the structure prediction of HLA-B with cumulative AA substitutions. Gray: the structure of 6UJB. Light blue: AlphaFold3-predicted structure with cumulative AA substitutions. Red: presented peptide (SPNGTIRNIL). (G) The structure comparison between the X-ray diffraction structure of human HLA-C (PDB: 6JTO) and the structure prediction of HLA-C with cumulative AA substitutions. Gray: the structure of 6JTO. Light blue: AlphaFold3-predicted structure with cumulative AA substitutions. Red: presented peptide (GADGVGKSAL).

HLA molecules showcase tumor-specific antigens derived from mutated or overexpressed proteins, potentially activating cytotoxic T cells to target malignant cells. The mechanism involves peptide binding within a specific groove on the HLA molecule, where the anchor residues of the peptide interact with complementary pockets in the HLA binding cleft, securing the peptide for recognition by T-cell receptors. Specifically, these complementary pockets are formed by three *α* domains in the HLA proteins [112]. AA substitutions within this binding *α* domians can significantly modify its physicochemical properties, such as size, charge, and hydrophobicity, thus altering the spectrum of peptides that can be effectively bound [113]. These AA substitutions may result in the inability to present immunogenic tumor antigens, facilitating immune evasion and cancer progression [114]. In this study, the majority of identified AA substitutions in MHC proteins are situated within the *α* domains. Specifically, HLA-A contains 9 AA substitutions within its *α* domains (Figure 7A), HLA-B has 13 substitutions (Figure 7B), HLA-C features 12 AA substitutions (Figure 7C), and HLA-DRB1 presents 18 AA substitutions (Figure 7D). AA substitutions within the *α* domains can alter the physicochemical properties of the key anchor pockets thereby radically reshaping the repertoire of peptides that can be effectively bound and presented to T-cells.

To continue quantificatively investigate the influence of the identified AA substitutions on the binding affinity between the MHC proteins and the presented peptides, we use the NetMHCpan to predict the affinity before and after the AA substitutions. The results of NetMHCpan show that the AA substitutions can significantly undermine the binding interaction between the MHC proteins and the presented peptides (Supplementary File 8). There are two strong binder are predicted between wild type HLA-A and the presented peptide VVVGAGGVGK, however, for the AA substituion modified HLA-A, there is no binder predicted (Figure 7E). The same observation applies to HLA-B, there are two binders, one is strong one is weak, predicted. But for AA substitution modified HLA-B, there is no predicted binder at all (Figure 7F). For HLA-C, the binding affinity is predicted by NetMHCpan, and the highest binding affinity is about top 10.9%, but the affinity of the AA substitution modified HLA-C is not predicted (Figure 7G).

Therefore, AA substitutions in MHC proteins hold significant importance in immunology and medicine, as these variations are key contributors to MHC polymorphism. This polymorphism, in turn, influences the diversity of the peptide presented to T cells and shapes an individual’s immune response to pathogens, vaccines, and cancer. In this paper, the AA substitutomic pipeline offers valuable insights into the structure-function relationship of MHC molecules, illustrating how specific residues within the binding groove determine peptide selectivity and binding affinity. This understanding is crucial for elucidating mechanisms of autoimmunity and immune evasion in cancer cells, thereby advancing the fields of personalized immunology and therapeutic interventions.

## 3 METHODS

### 3.1 Proteomic datasets and data preprocessing

In this study, proteomic data generated by CPTAC were downloaded from the Proteomic Data Commons [115]. The exploited data included 537 patients from five cancer cohorts: 11 TMT-11 plex experiments from 99 patients with GBM (PDC000204) [22], 20 TMT-11 plex experiments from 110 patients with HNSCC (PDC000221) [23], 25 TMT-10 plex experiments from 110 patients with LUAD (PDC000153) [24], 22 TMT-11 plex experiments from 108 patients with LSCC (PDC000234) [25], and 23 TMT-10 plex experiments from 110 patients with ccRCC (PDC000127) [26]. Besides, these additional cohorts for generalizability are sourced from ProteomeXchange [116] by the following data identifiers: PXD047816 (GBM) [117], PXD033898 (HNSCC) [118], PXD054681 (LUAD) [119], PXD035259 (LSCC) [120], and PXD012665 (ccRCC) [121]. The datasets enriched with cancer drug target proteins are sourced from ProteomeXchange, utilizing the following data identifiers: PXD000788 [91], PXD016465, PXD016505, and PXD016463 [92]. Additionally, the data identifiers for the MHC protein-enriched datasets available on ProteomeXchange are PXD003790 [110] and PXD000394 [111]. All of the non-empty MS2 spectra were extracted and converted to mgf files using ProteomeWizard MSConvert [122].

### 3.2 Analysis of proteomic data

The human protein FASTA file was downloaded from UniProt [28] (downloaded January 2024). PIPI-C was used as the open search method with the following parameters: the MS1 tolerance was 10 ppm; the MS2 tolerance was 0.01 Da; the max missed cleavage sites was 2; the enzyme was Trypsin; the peptide length was from 6 to 50; the fixed modifications were carbamidomethyl on cysteine and TMT labeling mass shift on lysine; modifications search range was ±250 Da; the variable modifications were from the UNIMOD [16], including TMT labeling mass shift on serine and n-term and all types of AA substitutions; isoleucine and leucine were treated as identical. Decoy proteins were added for FDR control with the target-decoy strategy. The remaining settings were kept as the default. Spectra were selected for the Comet search if PIPI-C identified more than two peptides with AA substitutions in them. The variable modifications for the Comet search were determined by PIPI-C’s search results for the spectra, and other search parameters of Comet were set as the same as those of PIPI-C. The validated PSMs, which were the same in the identification results of PIPI-C and Comet, were kept for further bioinformatic analysis.

### 3.3 Details of proteomic quantification

In the quantification step, we only kept the PSMs that were detected with TMT labeling mass shift. And the peptides with multiple PSMs were kept as quantifiable peptides. The intensities of reporter ions for each PSM were obtained from the initial MS2 spectra. Every sample was assumed to encompass a collection of unregulated peptides that had comparable abundances to those of the common reference (CR) sample. Therefore, in the normalized sample, these unregulated peptides should demonstrate a log TMT ratio that is centered at zero. As a result, we implemented a two-component normalization operation for the intensities of the TMT labels from the CR intensity [25].

Log ratios at the PSM level were computed as the average log values across patients within the corresponding PSM. Peptides sharing identical modification patterns were clustered into a UPSP. The average value of all PSM-level TMT ratios in each UPSP was used as the UPSP ratio. Each cancer cohort was composed of a set of distinct experiments. Therefore, the distinct average log ratios for each UPSP in each experiment were calculated. And we only selected those UPSPs that appeared in over 3/5 of the experiments for further analysis and quantification. One-sample t-test and BH-FDR correction were employed to determine the p-value and q-value for each UPSP, respectively.

### 3.4 GO and KEGG enrichment analysis

Proteins that had quantified (in quantification results, q-value*<*0.05 and fold change>3 or fold change*<*1/3) UPSPs with AA substitutions modifications were subjected to GO enrichment and KEGG enrichment analyses, where the background was the human genome database. Two-tailed Fisher’s exact test and BH-FDR correction were used to measure the statistical significance of the enrichment analyses. The visualizations in this study only included the enrichment terms with q-values less than 0.01 in GO and KEGG analysis.

The script for GO enrichment analysis was developed using GOATOOLS [123] by Python, while the KEGG pathway enrichment analysis was conducted using GSEApy [124] by Python.

### 3.5 Terminology and construction of protein-protein interaction network

We incorporated PPI data sourced from the STRING database [125], which aggregates interaction information from various sources such as text mining, experiments, databases, co-expression, neighborhood, gene fusion, and co-occurrence. The PPI network was visualized using Cytoscape (version 3.10.2) [126]. In this network representation, each pair of interacting proteins and the link between them were respectively denoted as nodes and edges.

### 3.6 Protein structure modeling, comparing and measuring

The protein structures were aligned by the Matchmaker function of ChimeraX (version 1.4) [127]. The structures were modeled by ChimeraX. The structure of FLNA Ig-like domains 3–5 was obtained by X-ray diffraction from Escherichia coli expression (PDB: 4M9P). The X-ray diffraction structure of vimentin (fragment 144–251) was from humans (PDB: 3UF1). The structure of human kidney fructose-bisphosphate ALDOB was obtained by cryo-electron microscopy (PDB: 8D44). AlphaFold3 built the predicted structures with cumulative AA substitutions using its web service. The provided sequences were the original sequences, in which the “I” was not replaced by “L”, and the AAs were substituted as the identified AA substitutions. When there was more than one possible AA substitution identified on the same site, we preferred the AA substitution with the higher regulation fold change. The predicted structure with the best pLDDT score was used for structure comparison and similarity measurement.

### 3.7 Quantification of binding affinity between the drug-target pairs

Binding affinities were predicted using CB-Dock2 [93]. The uploaded structures included both the wild-type drug targets from the PDB database and the AlphaFold3 predicted structures with AA substitutions. All proteins were uploaded in PDB format. The ligand used in CB-Dock2 was obtained from the DrugBank database [128], also in PDB format. For comparing changes in binding affinity, we selected the results that shared the same docking size and selected the results with the lowest Vina score. All other settings were maintained at their default values in CB-Dock2.

### 3.8 MHC binding affinity prediction

The predictions for MHC binding affinity and the count of MHC binders were generated using NetMHCpan 4.2 [109]. Both the wild-type sequences and the sequences modified by AA substitutions were submitted in fasta format to the web-based interface of NetMHCpan. A peptide length of 14 mer was chosen, and the representative from the HLA supertype was selected for the predictions. All other settings were maintained at their default values, with the binder threshold set at 2%, indicating that peptides predicted to fall within the top 2% of binding affinities are classified as predicted binders. The binders with the top 0.5% of binding affinities were regarded as strong binders.

To quantify the impact of AA substitutions in MHC proteins, we gathered the presented peptide sequences from the PDB database, using the same identifiers as those in Figure 7E-G for HLA-A, HLA-B, and HLA-C, respectively. We then entered the wild-type protein sequences along with the AA substitution-modified sequences as custom MHC protein sequences in NetMHCpan 4.2 to predict and compare their binding affinities.

## 4 DISCUSSION

In this study, AA substitution-profiling is performed based on the mass spectrometry-based proteomic search results. CPTAC provides proteomic datasets from five cancer types: GBM, HNSCC, LUAD, LSCC, and ccRCC. These five cancers are representative and located in different human body tissues. Therefore, the AA substitutomics analysis of these five cancers helps explore pervasive AA substitution patterns among different types of cancers. As for the proteomic search method, we implement a novel search pipeline in this paper, which balances the search range and search accuracy. To the best of our knowledge, empowered by PIPI-C this pipeline is the first one to cover all types of AA substitutions without the numerical limitation of modifications in each PSM. Although PIPI-C’s performance surpasses all other competitors through several rigorous tests, especially the test of identifying multiple AA substitutions in one peptide, the search results of PIPI-C are not guaranteed to be error-free. Therefore, to make the bioinformatic results as correct as possible, we use Comet as the validation step, following the PIPI-C search step. By incorporating the two search methods in our pipeline, the search range and search accuracy of identification results are balanced.

We identify consistent AA substitution patterns across five cancer cohorts, with E>M representing the most prevalent substitution among all E>X substitutions, and P>V emerging as the most frequent substitution within P>X substitutions. These patterns are consistently observed across both five CPTAC cohorts. Given that translation fidelity is a critical determinant of gene expression precision, and ribosomal errors during mRNA-to-protein translation occur stochastically at an estimated rate of 10^−4^ [129], we hypothesize that the prevalence of E>M and P>V substitutions may stem from two interrelated factors: 1. heightened metabolic reliance on methionine[130] and valine [131, 132] in cancer cells, and 2. elevated expression of methionyl-tRNA [133] and valyl-tRNA synthetase [134] in tumors. The consistent association of these high-frequency substitutions with cancer suggests their potential as putative therapeutic targets.

We investigate the protein-level regulatory effects of AA substitutions through quantitative proteomic analysis. By comparing significantly regulated AA substitutions with documented variants in UniProt [28], we demonstrate the biological relevance of our findings, with 13% overlap validating prior observations (Supplementary File 3). To trace the origin of these substitutions, we further compare our results against OncoDB [37], a comprehensive database of DNA/mRNA mutation-derived AA substitutions. Notably, most regulated AA substitutions identified in our study are absent from OncoDB (Supplementary File 4), suggesting they arise during protein synthesis rather than from genetic mutations. While conventional genomic and transcriptomic approaches primarily detect substitutions of genetic origin, proteomic methods have been limited by the lack of comprehensive open search techniques for large-scale modification detection. Our study addresses this gap by developing a robust AA substitutomics pipeline that enables high-throughput identification of protein synthesis-derived substitutions. This innovative approach not only facilitates the exploration of cancer biology but also holds substantial potential for investigating protein-level variations across diverse disease contexts, opening new avenues for AA substitutomics discovery.

We present a novel framework for examining and clarifying the mechanisms of drug and immune escape in cancer cells through the perspective of AA substitutomics. Nonetheless, our investigation offers merely one potential viewpoint for therapeutic exploration in cancer and cannot encompass all dimensions of cancer biology. Additionally, although AA substitutions are significant contributors to drug escape, an exclusive focus on this mechanism offers an incomplete and overly simplistic understanding of escaping. This perspective may neglect vital alternative pathways, including epigenetic alterations, metabolic reprogramming, or the upregulation of efflux pumps, which can impart escape independent of AA substitutions in the drug’s target protein [135]. Therefore, it is essential to conduct comprehensive, integrative research designed to address the multifaceted mechanisms of escape. Furthermore, a reduced MHC binding affinity in AA substitution-modified sequences does not definitively elucidate the immune escape mechanism. Immune evasion can also arise from mechanisms such as T cell receptor escape. While these MHC binding predictions serve as an important preliminary filter, it is crucial to validate potential neoantigens experimentally to confirm their ability to elicit an immune response [136].

In this study, we identify several cancer-associated proteins exhibiting multiple significantly regulated AA substitutions. While these proteins are biologically relevant to cancer progression, we acknowledge several important limitations. First, clinical sample accessibility and experimental constraints prevent immediate wet-lab validation of our findings. Second, the vast combinatorial space of possible AA substitutions, where any residue may be replaced by any other AA, exceeds current biological annotation in the literature, making causal interpretation challenging. Besides, we cannot presently determine whether these AA substitutions represent drivers of oncogenesis or secondary effects of tumor biology, which is a crucial question for future investigation. Furthermore, the inherent limitations of mass spectrometry analysis must be considered. The technique provides only a temporal snapshot of proteomic states, capturing population-averaged data rather than single-molecule resolution. Consequently, identified AA substitutions likely represent aggregate modifications distributed across multiple protein molecules rather than co-occurring on individual proteins. Our structural visualizations, therefore, depict cumulative substitution patterns and their composite structural impacts. Developing methods to resolve these substitutions at the single-molecule resolution remains an important challenge for the field, as does understanding the functional consequences of specific substitution combinations within individual protein molecules. Therefore, to fully realize the potential of AA substitutomics, a concerted effort from the broader community is essential.

## 5 CONCLUSION

AA substitutions are crucial for regulating cellular activities in normal and cancer cells, impacting complex signaling and physiology. Nonetheless, prior studies have so far scrutinized a restricted number of AA substitutions, while efficient open-search peptide identification tools have been lacking. To fill the research gap in profiling AA substitutions in cancer analysis, we used PIPI-C [17] to analyze all types of AA substitutions across diverse cancers. Among the identified AA substitutions and substitution sites, several have been documented as resulting from DNA or mRNA mutations in OncoDB [37], while the majority of the identified AA substitutions remain unrecorded. For some of the unrecorded AA substitutions, we have found support and evidence from proteomic studies and protein structure studies, which implies the confidence of our AA substitutions profile pipeline.

For the first time, this AA substitutomics study systematically characterizes multiple AA substitutions in cancer-associated proteins, with proteomic-level validation and structural analyses independent of genomic/transcriptomic data. From the five cancer cohorts, we identify two clinically relevant substitutions in HBB (F43S and E91D), previously confirmed by proteomic studies to significantly affect protein stability and functional activity. Structural investigations reveal a P584T substitution in FLNA domain 4, where the hydrophobic-to-hydrophilic transition (P>T) may perturb local protein interactions. In ALDOB, we observe an A175N substitution predicted to alter energy metabolism and molecular stability in cancer cells. Additionally, we noted evidence of alterations in protein structure and a reduction in drug-targt binding affinity and MHC-peptide binding affinity, which may contribute to investigate drug and immune escape in cancer cells. While literature supports some identified substitutions, others represent novel findings requiring further validation. The incomplete mechanistic understanding of these undocumented substitutions underscores the critical need for expanded proteomic investigations to AA substitutomics analysis and elucidate the precise roles of AA substitutions in oncogenic processes.

In summary, AA substitutions play a critical role in modulating protein function and tumor pathophysiology. Leveraging the PIPI-C framework, we present an approach for systematically investigating cancer-associated AA substitutions across diverse cancer types and contribute to establishing an AA substitutomics analysis pipeline. The high reproducibility of these substitutions within matched cancer cohorts underscores their generalizability within the cancer proteome, independent of tumor type. Functional validation through comprehensive literature integration demonstrates strong concordance across multiple independent studies, reinforcing the biological relevance of our findings. This study establishes the first AA substitutomics landscape in cancer biology, revealing the potential of AA substitutions as diagnostic biomarkers while providing a foundation for future mechanistic investigations in cancer biology.

## 6 DATA AVAILABILITY AND SUPPLEMENTARY FILES

The results of five CPTAC cohorts are available on Zenodo, with record number 13383150.

The search results of drug-target pair proteins are available on Zenodo, with record number 17669590.

The search results of MHC proteins are available on Zenodo, with record number 17669609. Supplementary File 1: The normalized ratios of each type of AA substitution across the five CPTAC cancer cohorts.

Supplementary File 2: The quantification results of the five CPTAC cohorts. It contains the AA substitution, peptide sequence, regulation ratio, and statistics of the results.

Supplementary File 3: The AA substitution details of all proteins in the CPTAC cohorts, and the AA substitutions that have been recorded in the UniProt database.

Supplementary File 4: The AA substitution details of the eight proteins that have multiple significantly modified AA, and the recorded sources of the AA substitution sites in the OncoDB database. Supplementary File 5: The AA substitution details of the drug target proteins.

Supplementary File 6: The output from NetMHCpan 4.2 of PLEC and LDHB includes the AA substitutions-modified versions and the wild-type versions.

Supplementary File 7: The AA substitution details of the MHC proteins.

Supplementary File 8: The result of NetMHCpan 4.2 predicted binders between MHC proteins and the presented peptides.

## 8 ACKNOWLEDGEMENTS

This work is partially supported by ITC grants MHP/033/20 and ITS/043/23, RGC grants 16102422, 16103621, R4012-18, C7015-23G, and T12-101/23N from the Hong Kong SAR government, Hetao Cooperation Zone Project HzQB-KCZYB-2020083, and internal grants 3030 009, BGF.001.2023, CSSET24SC01, Z1056 and OKT26EG06 from HKUST. We appreciate Tania Wilmshurst’s help in proofreading the manuscript. We thank the Generic Diagramming Platform (https://biogdp.com/) for providing the diagram drawing tools.

