## supplementary note for "Amino acid substitutomics: profiling amino acid substitutions at proteomic scale unveils biological implication and escape mechanism in cancer"

### for

[†] The authors contribute equally to this paper

### 1 Supplementary Note 1: Implementation and performance of PIPI-C

PIPI-C [1] formulates an optimization framework that addresses the challenge of identifying peptides with multiple PTMs from a combinatorial perspective, building upon the foundational concepts established by PIPI [2] and PIPI2 [3]. In the identification process, PIPI-C extracts short AA segments as tags from the MS2 spectra based on the relevant peaks. These tags are then employed to retrieve candidate sequences from an FM-indexed protein database [4] using a fuzzy and bidirectional matching technique. The mass difference between the peptide sequence and the precursor ion of the MS2 spectrum is interpreted as the total mass of potential PTMs. This information is utilized to construct mixed integer linear programming (MILP) models, which can be solved using Gurobi [5]. Subsequently, peptide candidates with characterized PTM patterns are assembled and reranked through a protein feedback module [6], allowing PIPI-C to generate a PSM list while controlling the FDR using a target-decoy strategy.

For tag extraction, PIPI-C combines tags of varying lengths to enhance the quality of the candidate list, leveraging the FM-index, which facilitates efficient querying of sequences of any length within the protein database. In the matching process, all target and decoy protein sequences are concatenated into a single long sequence to create the original FM-indexed database (o-FMDB). However, it is important to note that even a single erroneous AA in a tag can lead to retrieval failures. To address this, a fuzzy matching strategy is employed, which records the matched AAs and outputs the longest successfully matched substring. Nevertheless, fuzzy matching alone is inadequate when erroneous AAs are located at the beginning of a tag, as this results in initial matching errors. To overcome this limitation, a bidirectional matching approach is utilized alongside an additional reversed FM-indexed database (r-FMDB) generated from the reversed long sequence. In this setup, we attempt to match the reversed tag against the r-FMDB, where the backward matching of the reversed tag corresponds to a forward matching of the original tag. By integrating this fuzzy and bidirectional matching strategy, we consistently capture the correct portion of a tag, regardless of whether the erroneous AA is positioned at the N-terminus or C-terminus, thereby enhancing candidate retrieval sensitivity. Subsequently, overlapping tags are merged into a single entity if their peaks align, while tags with lower intensity are discarded as noise. Finally, peptide backbone candidates are ranked based on the cumulative intensities of the identified tags. This aggregation of evidence from multiple tags supporting the same peptide backbone sequence effectively minimizes redundant evaluations of individual tags.

Given a candidate pair consisting of a peptide backbone sequence and associated tags, the protein sequence can be divided into three sections: a N-terminal section, a C-terminal section, and a gap section defined by the tags. The mass shift within each section is determined based on the peak locations of the tags, calculated as the mass difference between the starting peak and the ending peak of adjacent tags. This indicates that the PTM patterns within these sections operate independently. Consequently, PIPI-C formulates the characterization of PTMs as an optimization problem by constructing an MILP model for each potential combination of PTMs (**Figure S1**). The objective of this model is to maximize the total intensity of the matched experimental peaks corresponding to all b- and y-ions. PIPI-C employs Gurobi to solve this optimization problem, ultimately yielding the optimized results for PTM characterization.

In addition to methodological advancements, PIPI-C has demonstrated high performance, surpassing two leading open-search methods, Open-pFind [7] and MODplus [8], in a series of validation experiments. To assess the precision and sensitivity of backbone identification and PTM characterization, seven simulated datasets with varying signal-to-noise ratios (SNRs), generated by AlphaPeptDeep [9], were utilized. Each MS2 spectrum predicted the intensities of b and y-ions using the protein Packet_Kmod as a template, sourced from ProteomeTools [10] through the ProteomeXchange Consortium [11] (data identifier PXD009449). Noise peaks were randomly added to the predicted signal peaks to produce complete MS2 spectra. Each dataset contained 124,248 MS2 spectra encompassing up to four PTMs from 18 selected modifications, distributed as follows: 135 spectra with 0 PTMs, 20,004 with 1 PTM, 92,357 with 2 PTMs, 11,263 with 3 PTMs, and 489 with 4 PTMs. Results indicated that as SNR decreased, both PIPI-C and Open-pFind maintained relatively stable performance, whereas MODplus exhibited a more rapid decline in effectiveness. PIPI-C particularly excelled with peptides featuring more than two PTMs. This performance was further validated using 21 synthetic datasets [10], which collectively included 1,023,540 MS2 spectra across 21 raw files, each synthesized with a maximum of one selected PTM on the specified AA. Despite Open-pFind's approach of characterizing PTMs through enumerating single PTM and 2-PTM combinations, PIPI-C consistently outperformed both Open-pFind and MODplus in these datasets. Moreover, PIPI-C's performance was validated using dimethyl-labeled soybean datasets, where peptides were labeled with either light or heavy dimethyl, ensuring no peptide contained both labels. This dataset comprised approximately 800,000 MS2 spectra from the ProteomeXchange Consortium (data identifier PXD034796). The performance evaluation relied on the number of PSMs with peptides fully labeled by either type of dimethyl label, revealing that PIPI-C identified 75% and 17% more fully labeled peptides than Open-pFind and MODplus, respectively. Additionally, performance on PTM combination identification was confirmed using the Petunia datasets (ProteomeXchange Consortium, data identifier PXD005470). In this search setting, enriched ubiquitination on lysine was treated as an unknown PTM, serving as a benchmark for identifying reliable PTM combinations. PIPI-C yielded more PSMs with PTM combinations, demonstrating superior capability in identifying multiple PTMs in real data. All validation results were filtered at a peptide-level FDR of 0.01, reinforcing that PIPI-C outperforms state-of-the-art open-search methods. PIPI-C is developed in Java, and its source code is accessible on https://bioinformatics.hkust.edu.hk/. All parameter files, result files, and simulation datasets are available on Zenodo at https://doi.org/10.5281/zenodo.14885715.

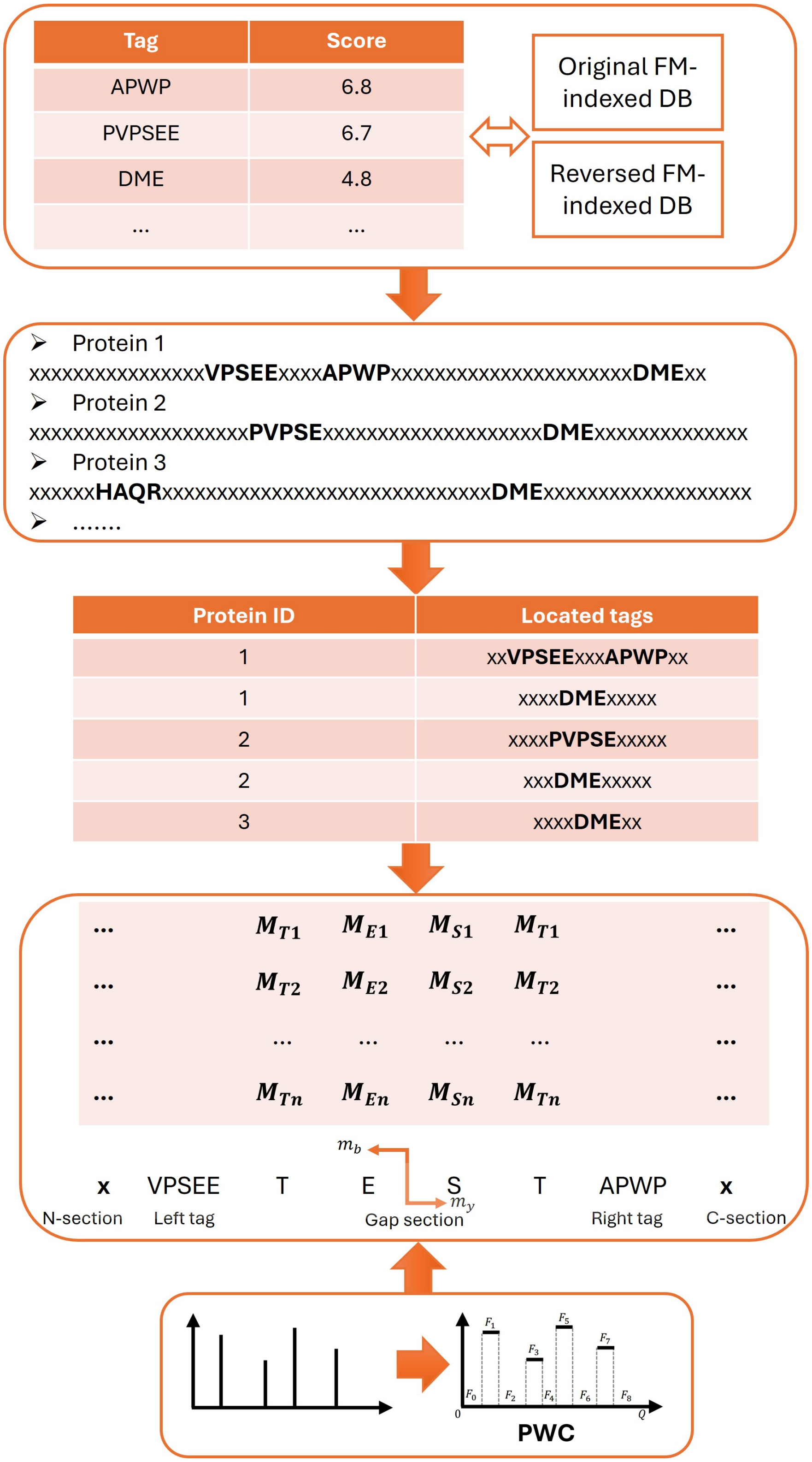
Tag
Score
APWP
6.8
PVPSEE
6.7
DME
4.8
...
...
Original FM-indexed DB
Reversed FM-indexed DB
Protein 1
xxxxxxxxxxxxxxxxVPSEExxxxAPWPxxxxxxxxxxxxxxxxxxxxxxxDMExx
Protein 2
xxxxxxxxxxxxxxxxxxxxPVPSExxxxxxxxxxxxxxxxxxxxxDMExxxxxxxxxxxxxxx
Protein 3
xxxxxxHAQRxxxxxxxxxxxxxxxxxxxxxxxxxxxxxxxDMExxxxxxxxxxxxxxxxxxxx
.......
Protein ID
Located tags
1
xxVPSEExxxAPWPxx
1
xxxxDMExxxxx
2
xxxxPVPSExxxxx
2
xxxDMExxxxx
3
xxxxDMExx
$M_{T1}$ $M_{E1}$ $M_{S1}$ $M_{T1}$
$M_{T2}$ $M_{E2}$ $M_{S2}$ $M_{T2}$
$M_{Tn}$ $M_{En}$ $M_{Sn}$ $M_{Tn}$
$m_b$
$m_y$
x
VPSEE
T
E
S
T
APWP
x
N-section
Left tag
Gap section
Right tag
C-section
$F_0$ $F_1$ $F_2$ $F_3$ $F_4$ $F_5$ $F_6$ $F_7$ $F_8$
0
$Q$
PWC

Figure S1: A schematic workflow for generating protein candidates is presented as follows. Initially, extracted tags are ranked based on their scores and used to retrieve protein candidates from FM-indexed databases. Next, the tags are aligned with the retrieved protein sequences using a fuzzy and bidirectional matching strategy. Subsequently, the protein sequences that contain the mapped tags are regarded as candidates and ranked according to the cumulative scores of the identified tags. Following this, the peaks of an MS2 spectrum are depicted as a piece-wise constant function. Finally, in both the N-terminal and C-terminal sections, amino acids are classified into fixed, optional, and infeasible categories to support the construction of the MILP model.

### 2 Supplementary Note 2: The experimental details of the false positive control test

In this study, we aim to profile various amino acid (AA) substitutions across five cancer cohorts. We design a series of experiments to evaluate and select the most suitable open search method for our research needs. The three candidate methods are PIPI-C [1], Open-pFind (version 3.2.0) [7], and MODplus (version 2.01) [8].

The first experiment is a false positive control test with simulation datasets. In this paper, we use AlphaPeptDeep [9] to generate simulation datasets. In each simulated MS2 spectrum, the intensities of b-ions and y-ions are predicted, and the noise peaks are randomly incorporated into the predicted peaks to produce the final MS2 spectra. In the false positive control test, we generate three simulation datasets with the average SNRs as 0.5, 1.0, and 2.0. The signal-to-noise ratio (SNR) is defined as:

$$\text{SNR} = \sum I_s^2 / \sum I_n^2. \tag{S1}$$

Where $I_s$ means the intensity of the signal peak and $I_n$ means the intensity of the noise peak. Each of the simulation datasets used for the false positive control test has 30,000 spectra. The parameters for AlphaPeptDeep to generate the simulation datasets in false positive control test are detailed in **Table S1**. The three methods are used with the following parameters: the MS1 tolerance: 10 ppm; the MS2 tolerance: 0.01 Da; the max cleavage site: 2; the enzyme: trypsin; the peptide length:

from 6 to 25; the fixed modification: carbamidomethyl on cysteine; the variable modifications: the UNIMOD [12] database including all types of AA substitutions and common post-translational modifications (PTMs); modifications search range: ±250 Da. The decoy proteins are added for FDR control with the target-decoy strategy. The rest of the settings are kept as the default. The human protein database is downloaded from UniProt [13] (January 2024). These search parameters are used in all testing experiments unless otherwise informed. To evaluate the performance of the false positive control test, since there are no AA substitutions or any modifications in the simulation dataset, the lower the ratio of identified AA substitutions, the better the performance. The heatmaps show the proportion of all kinds of AA substitutions (**Figure S2**). The proportion of each AA substitution is normalized by the number of the original amino acids in the result peptide spectra matches (PSMs):

$$\text{ratio} = \frac{\text{number of identified AA1>AA2 substitutions}}{\text{number of AA1 in the results}}. \tag{S2}$$

Table S1: Parameters used in AlphaPeptDeep for the simulated datasets in the false positive control test.

| | |
|---|---|
| **Enzyme** | trypsin |
| **Instrument** | Lumos |
| **Precursor charge** | 2-4 |
| **Peptide length** | 6-25 |
| **Max missed cleavages** | 2 |
| **Fixed modification** | Carbamidomethyl@C |
| **Num variable modifications** | 0 |

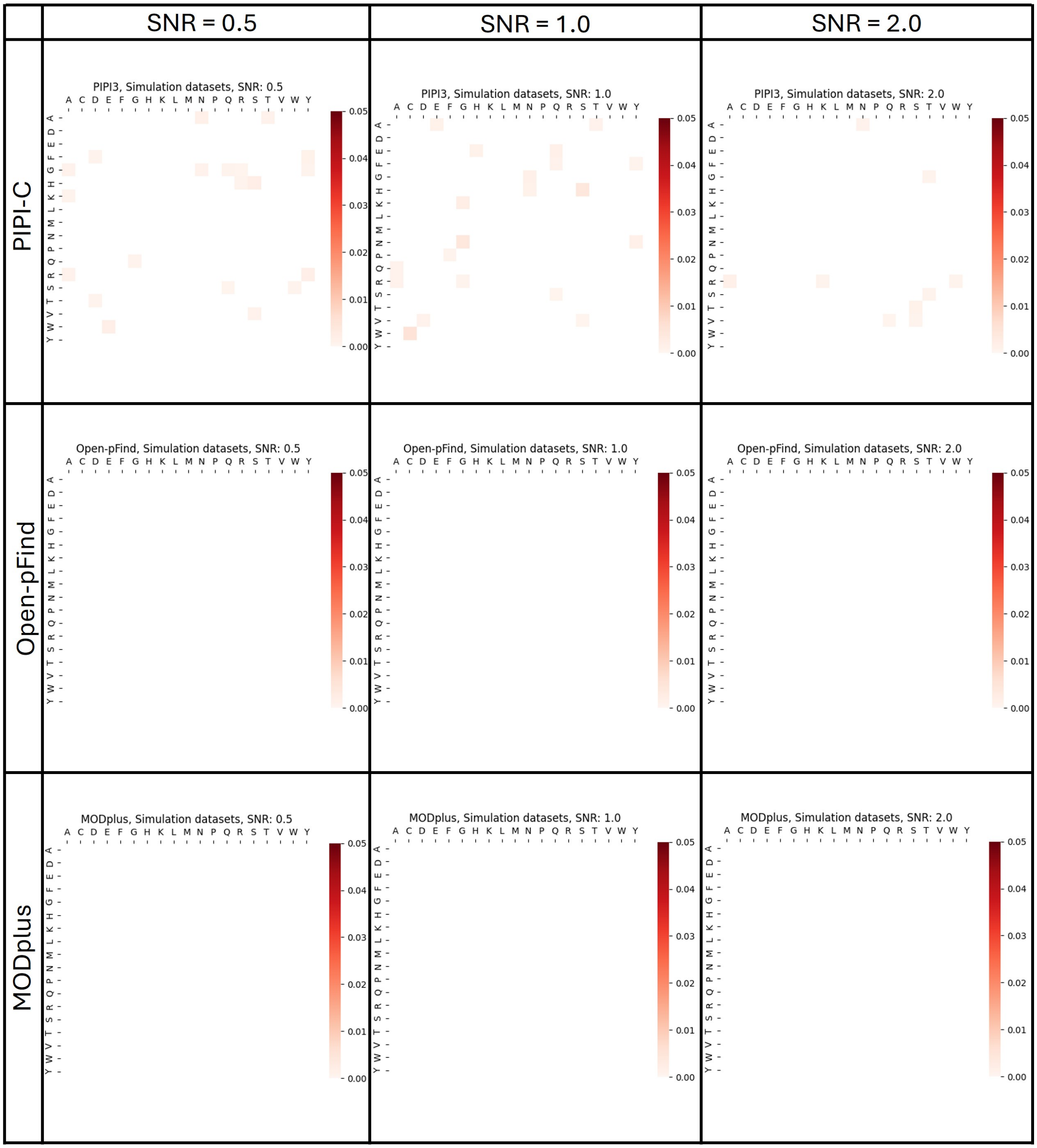


Figure S2: Results of PIPI-C, Open-pFind, and MODplus on the simulation datasets without AA substitutions. The spectra number of each dataset is 30,000.

From **Figure S2**, we can find that Open-pFind and MODplus detect no AA substitutions, while PIPI-C achieves a maximum ratio of approximately only 0.3% when the SNR is set to 1.0. The results of the false positive control test using the simulated datasets indicate a comparable performance among the three methods.

To conduct a more stringent false positive control test, we utilize 21 synthetic datasets obtained from ProteomeTools [10], which are sourced from the ProteomeXchange Consortium [11] with the data identifier PXD009449. The raw files are converted to the mgf format using ProteomeWizard MSConvert [14]. These datasets consist of 21 raw files containing a total of 1,023,540 MS2 spectra. In each raw file, the MS2 spectra are generated with at most one PTM on the specific amino acid. The details of the selected PTM and the modified residue are shown in **Table S2**. Given that there are no AA substitutions in the synthetic datasets, a lower ratio of identified AA substitutions indicates better performance. The normalized ratio of each type of AA substitution from the 21 datasets is illustrated in **Figure S3**.

Table S2: Information of the synthetic datasets in false positive control test.

| ID | Amino acid | Modification name | Monoisotopic mass | Number of MS2 |
|---|---|---|---|---|
| 1 | Lysine | Formylation | 27.995 | 51861 |
| 2 | Lysine | Acetylation | 42.010 | 50511 |
| 3 | Tyrosine | Phosphorylation | 79.966 | 55131 |
| 4 | Lysine | Methylation | 14.016 | 49086 |
| 5 | Lysine | Biotinylation | 226.078 | 43572 |
| 6 | Lysine | Butyrylation | 70.042 | 48573 |
| 7 | Lysine | Crotonylation | 68.026 | 49398 |
| 8 | Lysine | Dimethylation | 28.031 | 47914 |
| 9 | Lysine | Malonylation | 86.000 | 50877 |
| 10 | Lysine | Succinylation | 100.016 | 49896 |
| 11 | Proline | Hydroxyproline | 15.995 | 46893 |
| 12 | Lysine | Glutarylation | 114.032 | 49197 |
| 13 | Lysine | GlyGlycylation | 114.043 | 47664 |
| 14 | Lysine | Hydroxyisobutylation | 86.037 | 49329 |
| 15 | Lysine | Propionylation | 56.026 | 49305 |
| 16 | Lysine | Trimetylation | 42.047 | 49296 |
| 17 | Arginine | Citrullination | 0.984 | 50087 |
| 18 | Arginine | Dimethylation_asym | 28.031 | 45183 |
| 19 | Arginine | Dimethylation_symm | 28.031 | 45666 |
| 20 | Arginine | Methylation | 14.016 | 45396 |
| 21 | Tyrosine | Nitrotyrosine | 44.985 | 48705 |

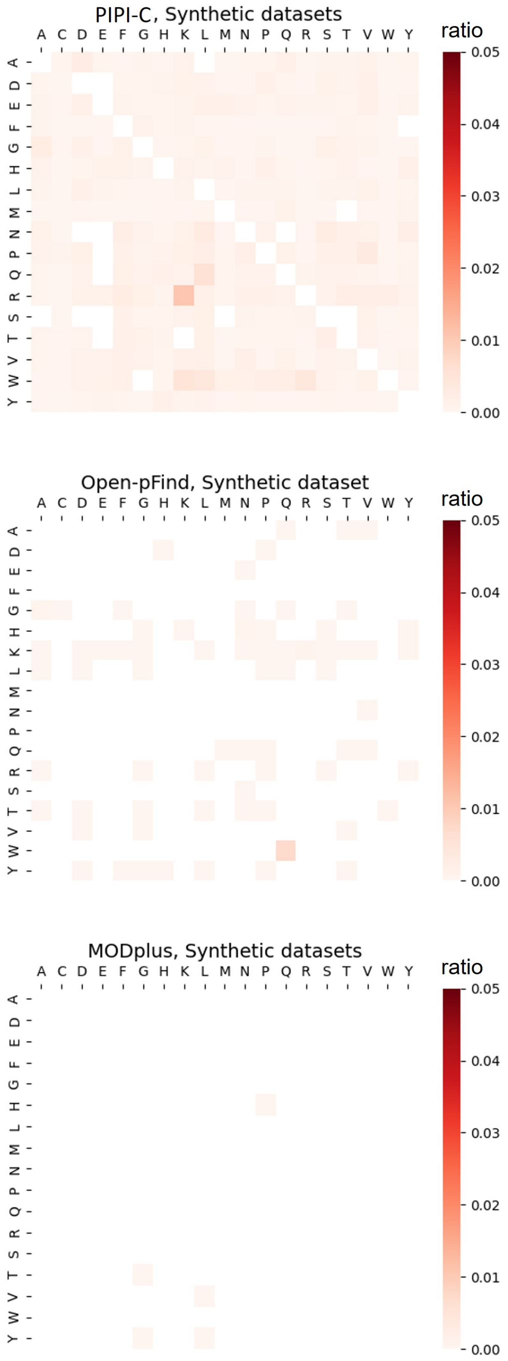


Figure S3: Results of PIPI-C, Open-pFind, and MODplus on the synthetic datasets without AA substitutions. The PSM number of the synthetic dataset is 1,023,540.

During our experiments with the synthetic datasets, we find the two substitutions S>D and T>E were frequently identified. Upon investigating the modification mass shifts of these two AA substitutions, we notice that both masses are equal to 27.995, which is identical to the mass of Formylation. Consequently, these two AA substitutions could be wrongly associated with Formylation, potentially arising from sample preparation steps. In mass spectrometry-based methods, it is not possible to distinguish such mass-identical AA substitutions and PTMs. Therefore, we find 20 modification patterns that have the same modification mass shift with AA substitutions (**Table S4**) and exclude these AA substitutions from the cancer data analysis. Based on **Figure S3**, it is observed that MODplus exhibits the most effective control over the false identification of AA substitutions, with a maximum false identification ratio of approximately 0.004% and a mean ratio less than $2 \times 10^{-7}$. Open-pFind follows as the second-best method, achieving a maximum false identification ratio of about 0.7% and a mean ratio of around $3 \times 10^{-5}$. PIPI-C shows a maximum false identification ratio of about 1% and a mean ratio of approximately $7 \times 10^{-4}$. In the results of PIPI-C and Open-pFind, only three and one AA substitutions, respectively, have a ratio exceeding 0.5%. Therefore, the false positive control abilities of the three methods on synthetic datasets are at the same level and satisfactory.

Based on the low identification rates discussed above, it can be concluded that the three methods exhibit comparable and satisfactory performance in the false positive control tests.

### 3 Supplementary Note 3: The experimental details of the AA substitutions identification sensitivity test.

In the test for sensitivity of detecting AA substitutions, we utilize AlphaPeptDeep to generate three simulation datasets with varying numbers of AA substitutions in a single peptide: one AA substitution per peptide (1-AASub dataset), two AA substitutions per peptide (2-AASub dataset), and three AA substitutions per peptide (3-AASub dataset). Given that carbamidomethyl is considered the fixed modification on cysteine and isoleucine and leucine are treated as leucine, there are a total of 324 types of AA substitutions in this study. Each simulation dataset consisted of 324 spectrum files, with each file containing 1000 spectra. Among these spectra, 5% include one type of AA substitution. The parameter configurations for AlphaPeptDeep are outlined in **Table S3**. The search parameters remain consistent with those employed in the false positive control test.

Since the ratio of spectra containing AA substitutions is known to be 5%, we utilize the absolute difference between the identified AA substitution ratio and 5% as the evaluation metric. A smaller difference indicates that the AA substitutions ratio closely aligns with 5%, signifying better sensitivity in detecting AA substitutions. The outcomes of the three methods across the three simulation datasets with varying numbers of AA substitutions per peptide are depicted in **Figure S4**. Upon analyzing the results of the sensitivity test, it is observed that Open-pFind exhibits slightly better performance in the 1-AASub simu dataset than the other methods. However, in scenarios involving multiple substitutions, PIPI-C surpasses Open-pFind and MODplus. Particularly, in settings with multiple AA substitutions, Open-pFind only correctly identifies a limited number of AA substitutions, falling short of meeting the desired criteria.

Table S3: Parameters used in AlphaPeptDeep for the simulated datasets in the test for sensitivity of detecting AA substitutions.

| | |
|---|---|
| **Enzyme** | trypsin |
| **Instrument** | Lumos |
| **Num variable modifications** | 1 or 2 or 3 |
| **Max missed cleavages** | 2 |
| **Precursor charge** | 2-4 |
| **Peptide length** | 6-25 |
| **Fixed modification** | Carbamidomethyl@C |
| **Variable modification** | 324 kinds of AA substitutions |

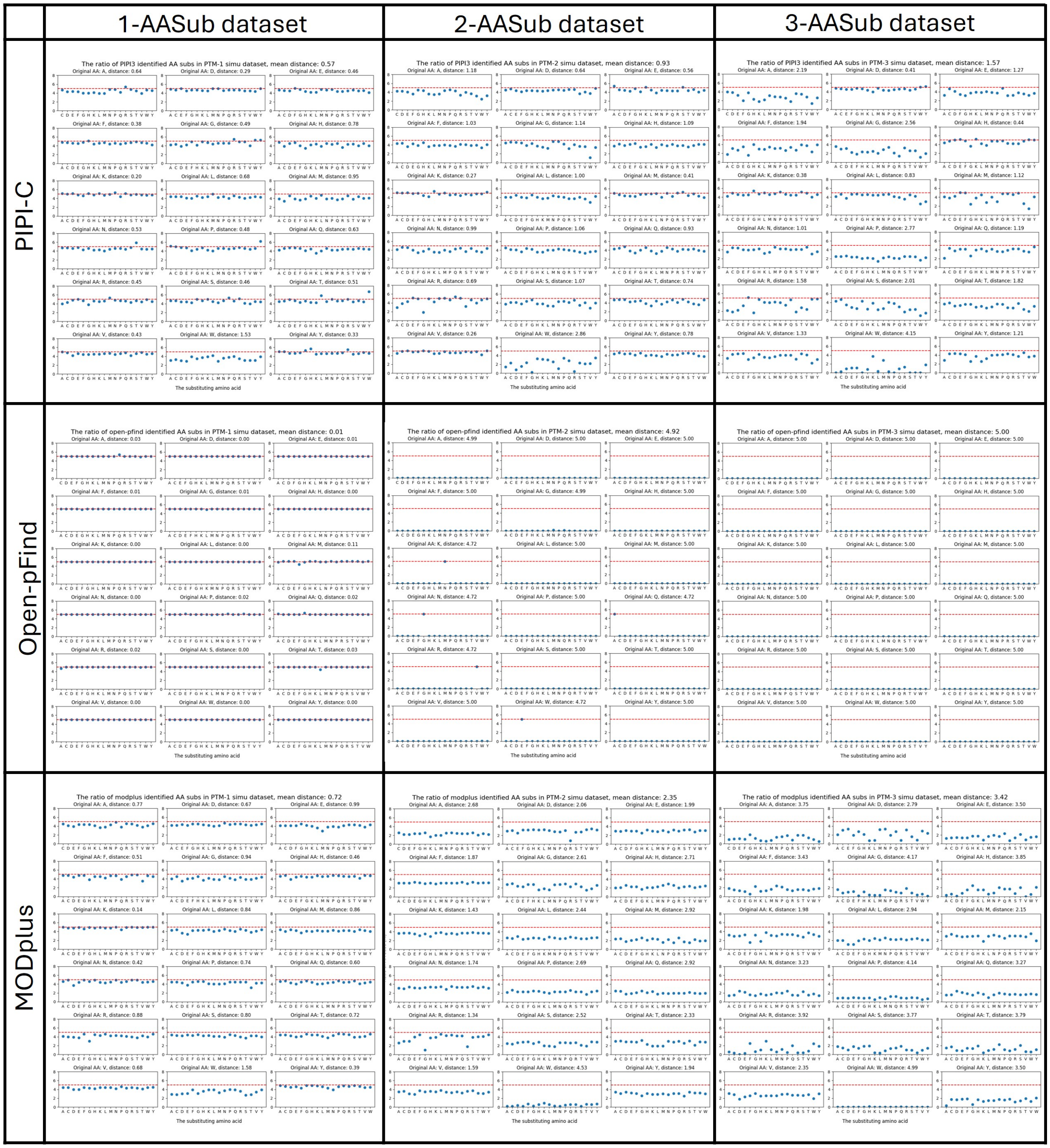


Figure S4: The results of PIPI-C, Open-pFind, and MODplus on the simulation datasets with 1–3 AA substitutions in each peptide. In each subplot, the x-axis is the substituting amino acid, the y-axis is the ratio. The original amino acid and the mean distance between the ratio and 5% are in the title of the subplots. The horizontal red dashed line is 5%. The spectra number of each dataset is 324,000.

### 4 Supplementary Note 4: The experimental details of the test for detecting multiple modifications.

In this paper, we aim to analyze the landscape and profile of all types of AA substitutions. Therefore, the ability to identify multiple AA substitutions in one peptide is crucial in method evaluation. Since there is no proteomic dataset with enriched AA substitutions, to fulfill this requirement, we use a proteomic dataset with serial enrichment of phosphorylation, ubiquitination, and acetylation [15]. The dataset is downloaded from the MassIVE Repository with the data identifier MSV000078509. After converting to MGF format, the dataset contains 612,348 spectra.

In **Figure S5**, we show the number of PSMs with different numbers of enriched PTMs in one peptide. The results from the three methods indicate that the three methods are comparable in identifying a single modification within a peptide. Additionally, when it comes to detecting multiple modifications in one peptide, PIPI-C demonstrates superior performance with many more identification results over both competing methods, with MODplus ranking as the second-best option.

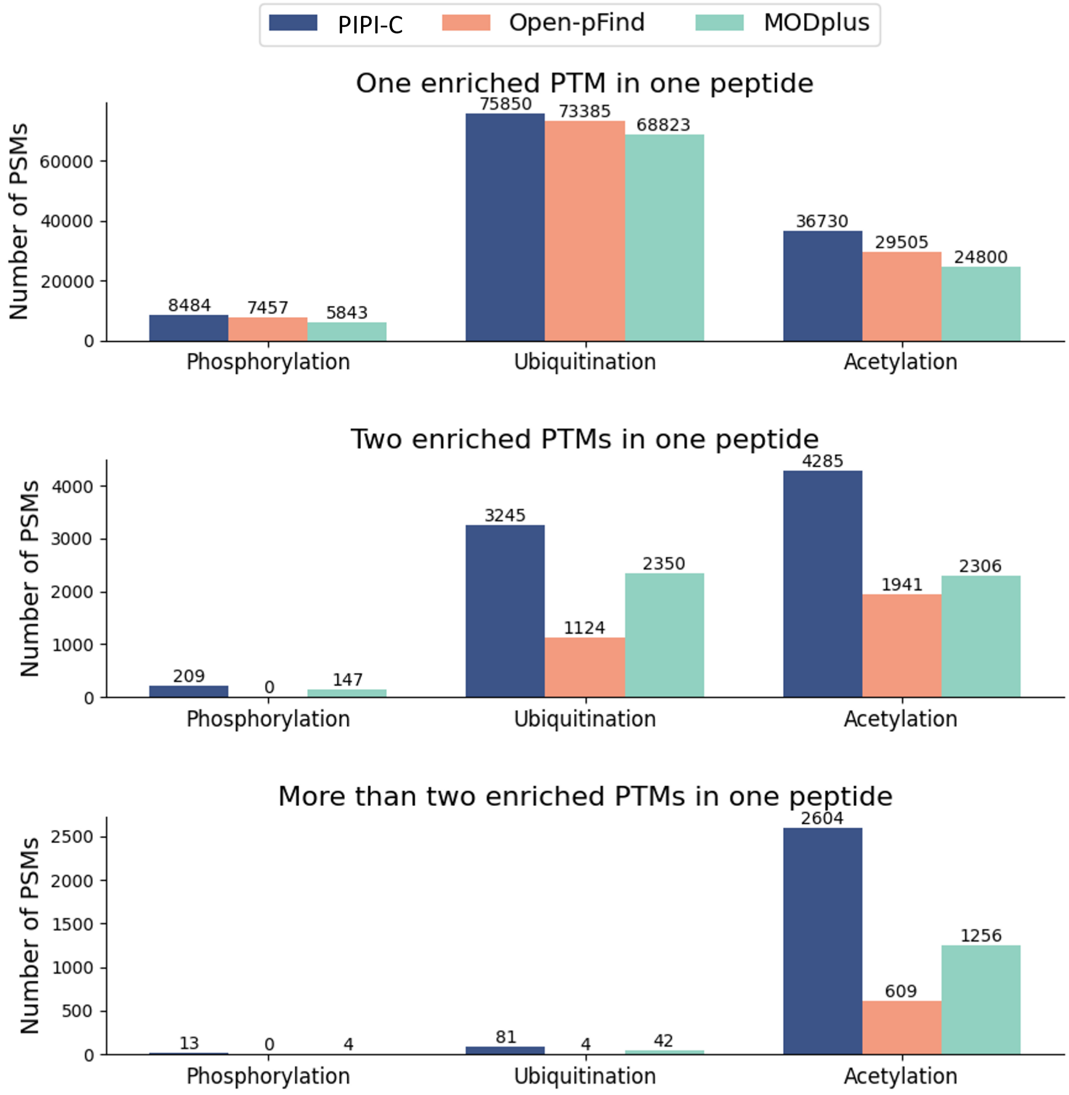


Figure S5: Number of PSMs with different numbers of enriched PTMs in one peptide from results of PIPI-C, Open-pFind, and MODplus on the serial PTM enrichment dataset. The dataset contains 612,348 spectra.

### 5 Supplementary Note 5: PIPI-C search result on proteomic data of mistranslating tRNA mutant and wild type.

In this section, we will showcase the power of PIPI-C in identifying AA substitutions on mistranslating tRNA mutant datasets, where the number of R>S substitution and P>A substitution is known to be more than in the wild-type (WT) dataset [16]. The dataset is downloaded from ProteomeXchange Consortium with the data identifier PXD025934 and the protein database of Saccharomyces cerevisiae (yeast) is downloaded from UniProt (October 2024). When growing yeast, two types of engineered tRNA are involved, therefore, the frequencies of R>S substitution and P>A substitution in the two mutant groups are higher than in the WT group

PIPI-C identifies 156,165 PSMs, 121,096 PSMs, and 152,371 PSMs for the WT group, R>S group, and P>A group, respectively. From **Figure S6**, we can find that PIPI-C successfully identifies more designed AA substitutions in each mutant group compared with the WT group and another mutant group. With this experiment, PIPI-C is proven to differentiate the mutant sample and the WT sample with different frequencies of AA substitutions. Such observation further encourages us to apply PIPI-C to the real cancer dataset and makes us more confident about PIPI-C's results.

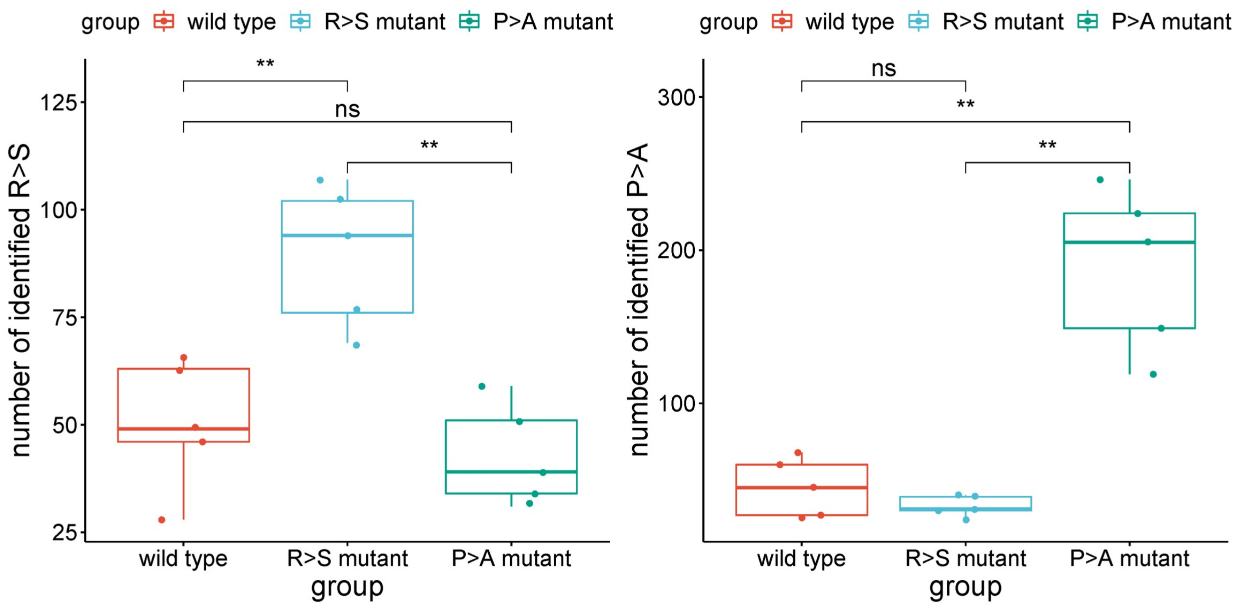


Figure S6: Boxplot showing the numbers of PIPI-C identified AA substitutions are significantly different between the mutant datasets and the WT datasets. Left: the number of identified R>S substitutions. Right: the number of identified P>A substitutions. T-test p-values are shown, **: p-value $< 0.01$, ns: p-value $> 0.05$.

### 6 Supplementary Note 5: Other supplementary materials.

Table S4: The ambiguous AA substitutions have the same mass shift as other modifications.

| Modification site | Monoisotopic mass | AA substitution name | Modification name |
|---|---|---|---|
| Aspartic acid | 14.016 | Asp>Glu | Methylation |
| Phenylalanine | 15.995 | Phe>Tyr | Oxidation |
| Methionine | -29.993 | Met>Thr | Met>Hse |
| Asparagine | 0.984 | Asn>Asp | Deamidated |
| Glutamine | 0.984 | Gln>Glu | Deamidated |
| Serine | -15.995 | Ser>Ala | Deoxydation |
| Serine | 14.016 | Ser>Thr | Methylation |
| Threonine | 27.047 | Thr>Lys | Ethylamino |
| Alanine | 42.047 | Ala>Xle | Trimethylation |
| Asparagine | 15.000 | Asn>Glu | Methylation+Deamidated |
| Asparagine | 14.016 | Asn>Gln | Methylation |
| Proline | 31.990 | Pro>Glu | Dioxidation |
| Serine | 27.995 | Ser>Asp | Formylation |
| Serine | 42.011 | Ser>Glu | Acetylation |
| Serine | 44.008 | Ser>Met | Delta:H(4)C(2)O(-1)S(1) |
| Threonine | 27.995 | Thr>Glu | Formylation |
| N-term | 14.016 | Asp>Glu<br>Ser>Thr<br>Val>Xle<br>Asn>Gln<br>Gly>Ala | Methylation |
| C-term | 14.016 | Asp>Glu<br>Ser>Thr<br>Val>Xle<br>Asn>Gln<br>Gly>Ala | Methylation |
| N-term | 27.995 | Ser>Asp<br>Thr>Glu | Formylation |
| N-term | 42.011 | Ser>Glu | Acetylation |

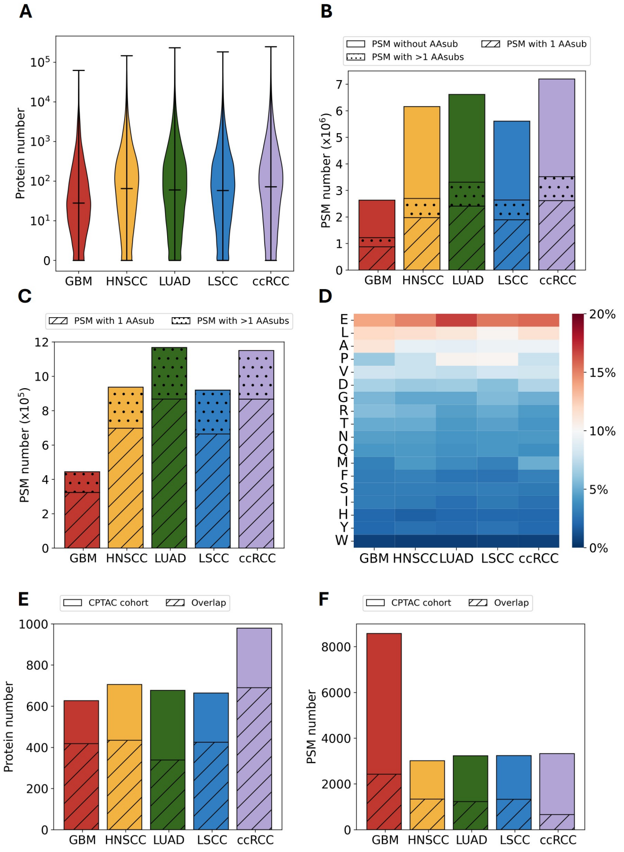


Figure S7: The search results of the AA substitutomic pipeline. (A) Violin plot showing the PIPI-C-identified protein number of each cancer cohort. The cohorts are GBM (dark red), HNSCC (yellow), LUAD (green), LSCC (blue), and ccRCC (purple).
(B) The PSMs in PIPI-C's identification results. Clean section: PSMs without any AA substitutions. Slashed section: PSMs with one AA substitution. Dotted section: PSMs with more than one AA substitution. The cohorts are GBM (dark red), HNSCC (yellow), LUAD (green), LSCC (blue), and ccRCC (purple).
(C) Bar plot showing the number of PSMs in Comet's validated identification results. Slashed section: PSMs with one AA substitution. Dotted section: PSMs with more than one AA substitution. The cohorts are GBM (dark red), HNSCC (yellow), LUAD (green), LSCC (blue), and ccRCC (purple).
(D) Normalized ratio of substituted AAs in each cohort.
(E) The overlap results of identified protein numbers. The cohorts are GBM (dark red), HNSCC (yellow), LUAD (green), LSCC (blue), and ccRCC (purple). Slashed section: identified proteins that overlap with the CPTAC cohort. The overlap ratios of proteins identified in additional cancer cohorts with the regulated proteins from the CPTAC cohorts range from 50% to 71%.
(F) The overlap results of PSM numbers. The cohorts are GBM (dark red), HNSCC (yellow), LUAD (green), LSCC (blue), and ccRCC (purple). Slashed section: identified peptides that overlap with the CPTAC cohort. The PSM-wise overlap ratios vary between 20% and 44%, with the ccRCC and GBM cohorts exhibiting the lowest overlap ratios at 20% and 28%, respectively.

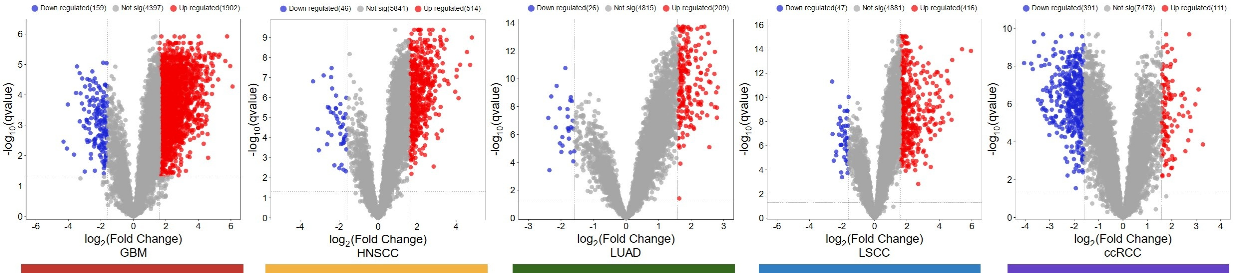


Figure S8: Volcano plots of quantification results based on the validated PSMs with modifications from five cohorts.

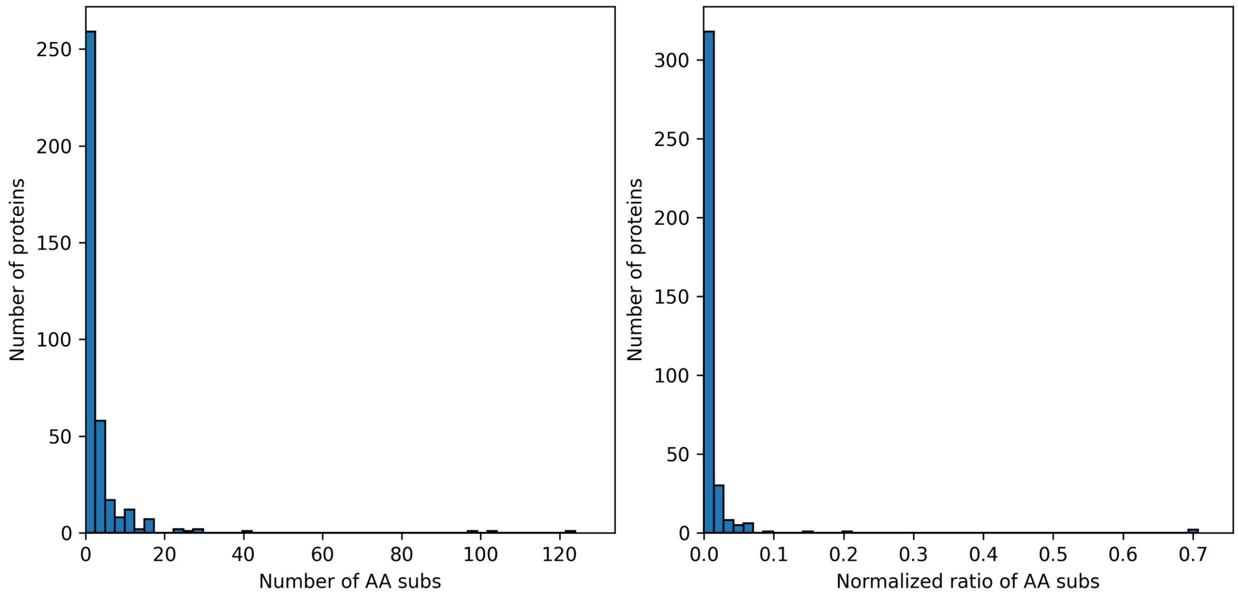


Figure S9: The histogram of the number of identified AA substitutions in proteins. Left: before normalization by the protein length. Right: after normalization by the protein length.

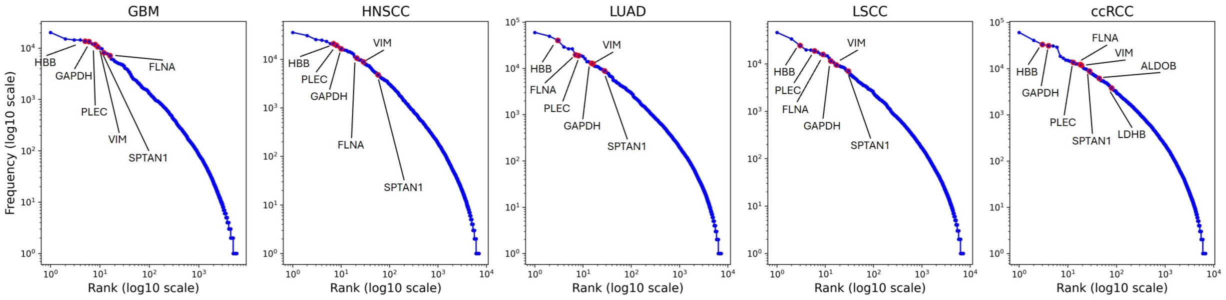


Figure S10: The abundance of the proteins that have multiple significantly regulated AA substitutions

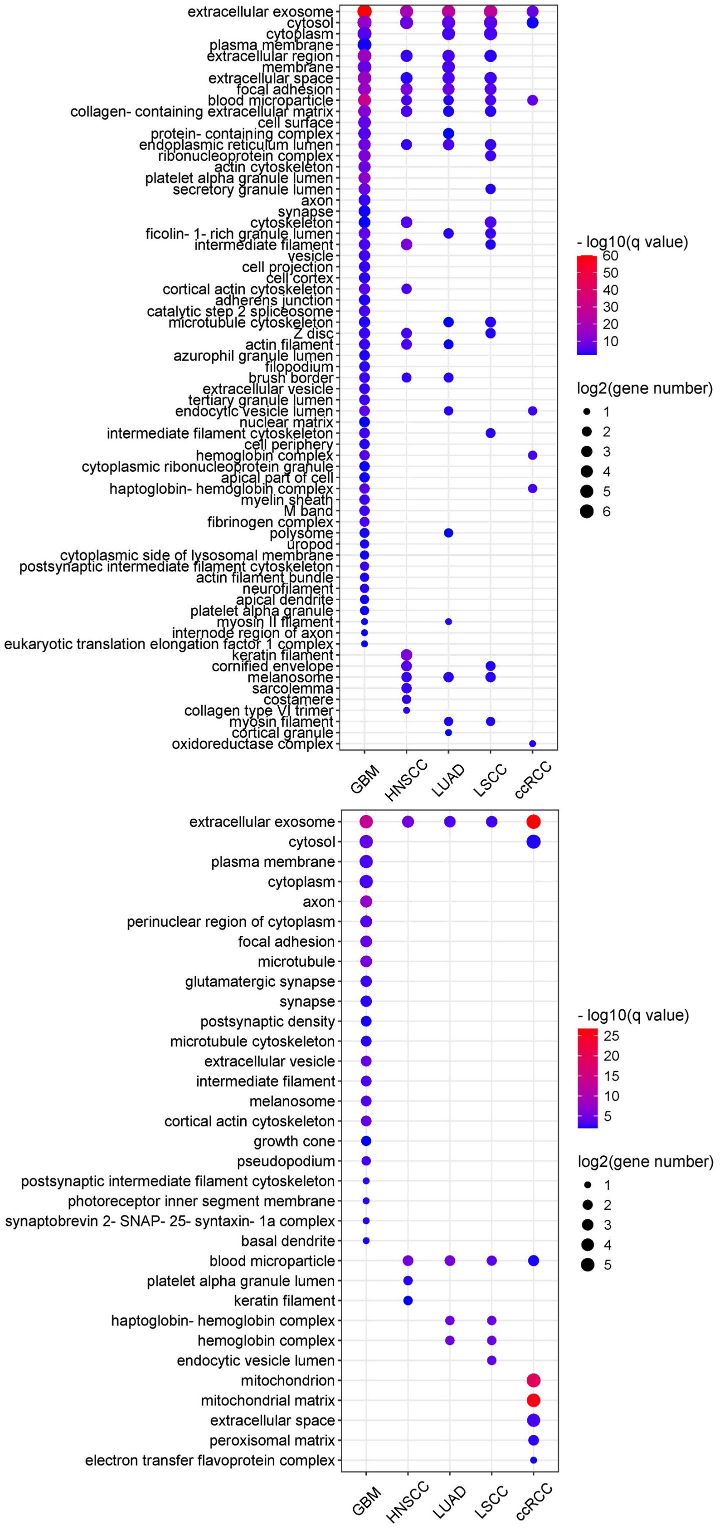


Figure S11: The GO cellular component enrichment results based on the significantly regulated UPSPs. Top: the cellular component enrichment based on up-regulated UPSPs. Bottom: the cellular component enrichment based on down-regulated UPSPs. The size and color of the bubble represent the number of genes enriched in each molecular function and their minus base 10 logarithm of q value, respectively.

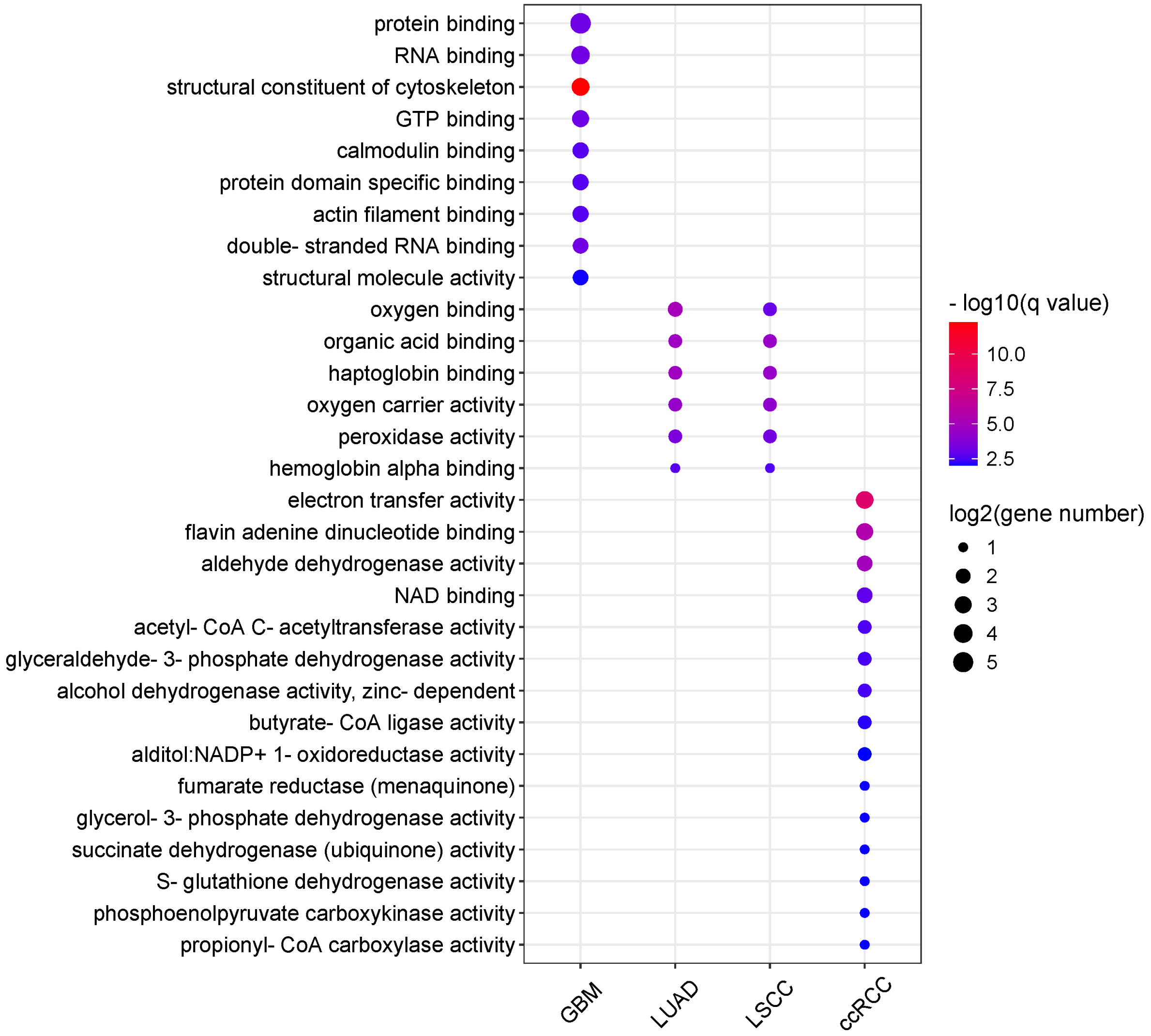


Figure S12: The GO molecular function enrichment results based on the down-regulated UPSPs. The size and color of the bubble represent the number of genes enriched in each molecular function and their minus base 10 logarithm of q value, respectively.

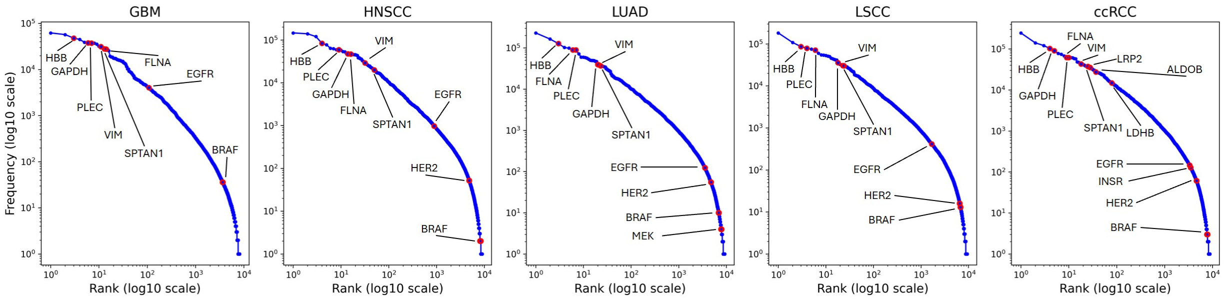


Figure S13: The abundance of the proteins that have multiple AA substitutions and the current cancer drug target proteins from the PIPI-C's identification results.

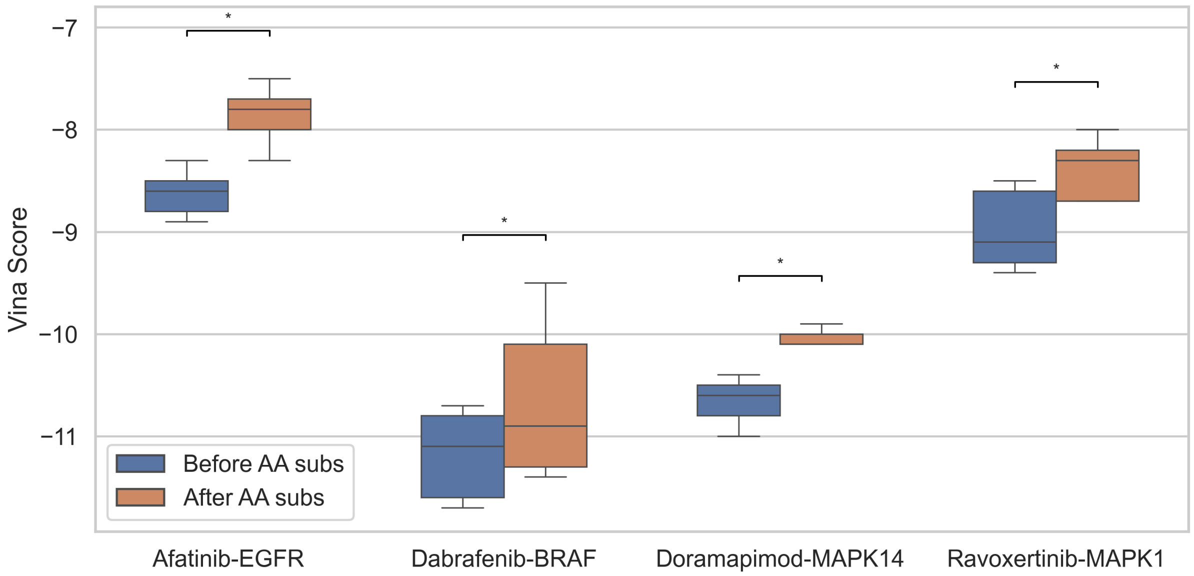


Figure S14: The change of Vina score change before and after the AA substitutions of the four drug-target pairs. T-test p-values are shown, ⋆: p-value $< 0.05$.

[10] D. P. Zolg, M. Wilhelm, T. Schmidt, G. Médard, J. Zerweck, T. Knaute, H. Wenschuh, U. Reimer, K. Schnatbaum, B. Kuster, *et al.*, "ProteomeTools: Systematic characterization

of 21 post-translational protein modifications by LC-MS/MS using synthetic peptides," *Mol. Cell. Proteomics*, pp. 1850–1863, 2018.
